# Eosinophils promote monocyte to macrophage differentiation and anti-bacterial immunity

**DOI:** 10.1101/2024.11.25.625164

**Authors:** Gillian J Wilson, Lily Koumbas Foley, Zuzanna Pocalun, Elise Pitmon, Ayumi Fukuoka, Kit M Lee, Lauren Fernandez, Heather Mathie, Robin Bartolini, John J. Cole, Gerard J Graham

## Abstract

Chemokine receptors control cell migration within the body. Here we reveal a novel interaction between eosinophils and monocytes in the bone marrow, indirectly controlled by the atypical chemokine receptor ACKR2. We demonstrate that ACKR2 maintains eosinophil levels within the bone marrow by scavenging CCL11. In the absence of ACKR2, elevated CCL11 leads to increased egress of eosinophils from the bone marrow into the bloodstream. As a result, eosinophil and monocyte interactions are reduced within the bone marrow niche, leading to changes in monocyte gene expression. Monocytes from ACKR2-/- mice are recruited to the tissues but are fundamentally altered in their ability to differentiate into macrophages, in the lung, peritoneal cavity and cavity wall. Bacterial elimination is impaired in ACKR2-/- mice during peritoneal infection. ACKR2 is therefore a key regulator of eosinophil-driven monocyte education in the bone marrow, required for full monocyte differentiation and macrophage function within the tissues.

## INTRODUCTION

Macrophages are key cells within the immune system, protecting the body by phagocytosing pathogens, helping to repair and remodel tissues after injury, and maintaining tissue homeostasis ^1^. Macrophages in the tissues can be broadly divided into resident or recruited cells. Resident macrophages are seeded embryonically and derive from yolk sac and foetal liver progenitors ^2^. Recruited/inflammatory macrophages are derived from bone marrow monocytes which differentiate into macrophages following recruitment into tissues ^3^.

Monocyte and macrophage movement within the body is largely orchestrated by members of the chemokine family ^4, 5^. Chemokines are a family of proteins characterised by a conserved cysteine motif and the large chemokine family is subdivided according to cysteine distribution into CC, CXC, XC and CX3C subfamilies. Chemokines act through G-protein-coupled receptors to direct cell migration ^6, 7^. Monocyte mobilisation and recruitment is predominantly regulated by the CC receptor 2 (CCR2), and CX3C receptor 1 (CX3CR1) ^8^. In addition to signalling chemokine receptors, there is a subfamily of atypical chemokine receptors (ACKRs), that lack classical signalling responses to their ligands ^7, 9^. ACKR2 is a promiscuous scavenger receptor which binds, internalises and targets inflammatory CC-chemokines for degradation within the cell ^6^. By modulating the availability of chemokines, ACKRs can regulate chemokine gradients and alter cell recruitment and activation. Together, signalling chemokine receptors and ACKRs coordinate leukocyte migration within the tissues ^6^.

We, and others, have demonstrated an essential role for ACKR2 in resolving CC-chemokine driven inflammatory responses and in controlling the development of inflammation dependent cancers ^10, 11, 12, 13, 14, 15^. In addition, we have demonstrated that ACKR2 plays key roles in development, scavenging chemokines in the placenta, embryonic skin, and the pubertal mammary gland ^16, 17, 18, 19^. In embryonic skin, ACKR2 is expressed by lymphatic endothelial cells (LECs) and scavenges CCL2 to control the proximity of CCR2-expressing macrophages to developing vessels ^18^. In the developing mammary gland, ACKR2 is expressed by fibroblasts and scavenges CCL7 to control CCR1-expressing macrophages and epithelial branching ^16, 17^.

Eosinophils are granulocytic immune cells derived from haematopoietic stem cells (HSC) in the bone marrow ^20^. They are classically associated with the immune response in allergy or parasitic infection, but more recently, their roles in maintaining tissue homeostasis and shaping development in the thymus have been identified ^20, 21^. Here we report a previously unknown role for eosinophils in educating monocytes within the bone marrow, indirectly controlled by the atypical chemokine receptor ACKR2. We demonstrate that in the absence of ACKR2 scavenging, the concentration of CCL11, a ligand shared with the dominant eosinophil chemokine receptor CCR3 ^22, 23^, is increased. We show that elevated eosinophil egress from the bone marrow to the blood in ACKR2-/- mice, results in reduced eosinophil/monocyte interactions within the bone marrow niche, and changes in monocyte gene expression. Monocytes from ACKR2-/- mice are recruited to the tissues but are fundamentally altered in their ability to differentiate into macrophages. Furthermore, in the absence of ACKR2, bacterial killing is impaired during peritoneal infection. Our data therefore demonstrate that ACKR2 is a key regulator of eosinophil-driven monocyte education in the bone marrow, which is required for full monocyte differentiation and ultimately macrophage function during infection.

## RESULTS

### Macrophage numbers are decreased in the peritoneal cavity, cavity wall and lungs of ACKR2-/- mice

In the peritoneal cavity, there are two main macrophage populations, small peritoneal macrophages (SPMs) which differentiate from CCR2 expressing Ly6C high monocytes, and resident large peritoneal macrophages (LPMs) ^24, 25, 26, 27^. These populations were identified using the flow cytometry gating strategy shown in Supplementary Figure 1. In the absence of ACKR2, the number of SPMs were reduced in the peritoneal fluid and the cavity wall of 8 week old female mice (Figure 1a,b). We saw the same reduction in female mice of different ages, 4 and 12 weeks, and in male 8 week old mice (Supplementary Figure 2). There was no effect on the number of resident LPMs in ACKR2-/- mice (Figure 1a,b). Upon depletion using clodronate liposomes, SPMs were replaced in WT mice but remained at lower levels in ACKR2-/- mice (Supplemental Figure 3). Together these data suggest that the SPM defect in the ACKR2-/- peritoneal cavity is stable and recapitulated following replenishment of the macrophage population.

**Figure 1:**
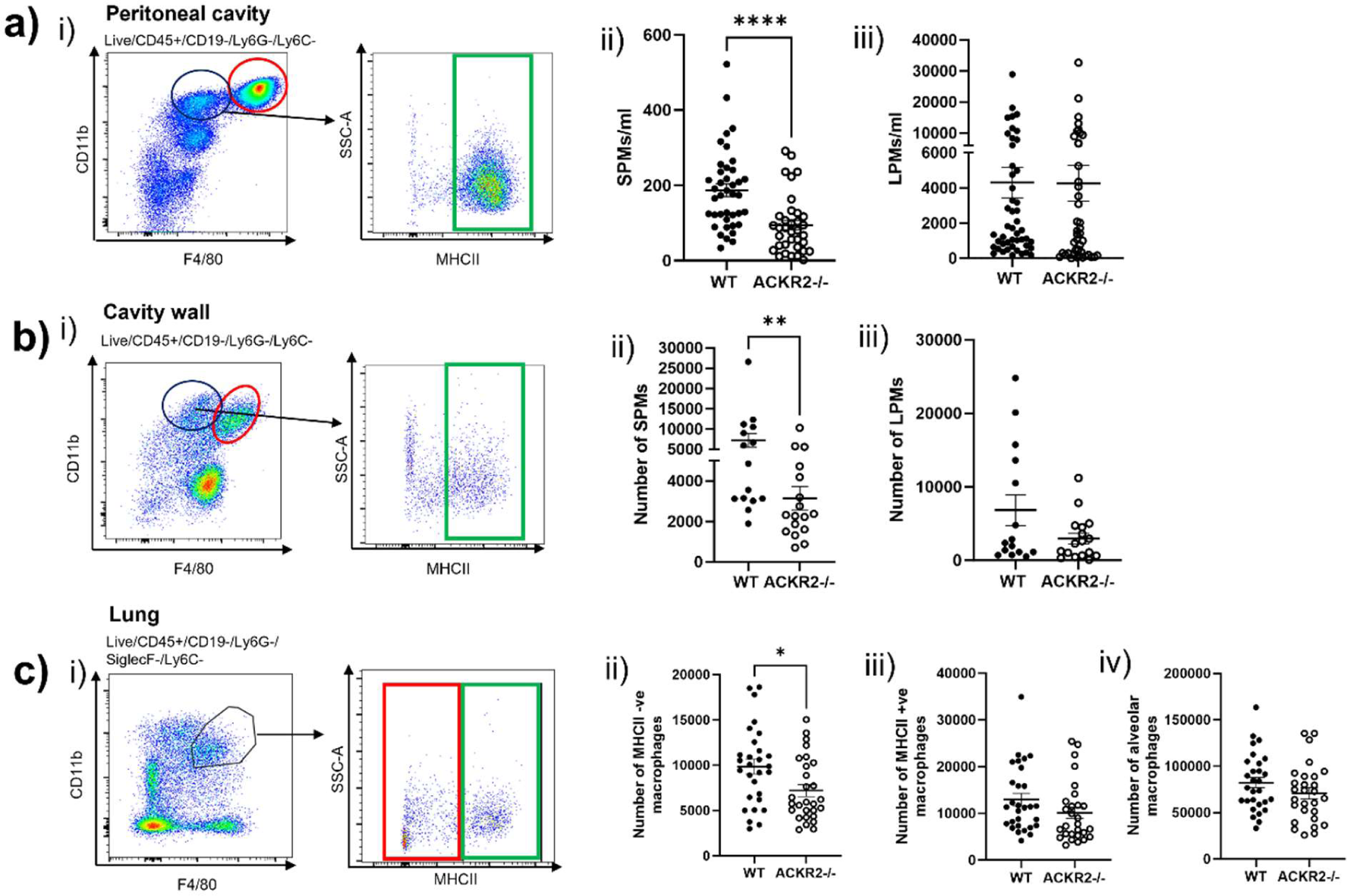
Macrophages are decreased in ACKR2-/- mice. **a) i)** Gating strategy to define large peritoneal macrophages (LPMs, circled in red) and small peritoneal macrophages (SPMs, green box) in the peritoneal cavity. Flow cytometry was carried out to determine the number of **ii)** SPMs, and **iii)** LPMs, WT *n*=40, ACKR2-/- *n*=36 in the peritoneal cavity. **b) i)** LPMs, circled in red, and SPMs, boxed in green, in the peritoneal cavity wall. The number of **ii)** SPMs and **iii)** LPMs, WT *n*=15, ACKR2-/- *n*=17, in the peritoneal wall. **c) i)** Gating of lung interstitial macrophages, MHCII positive (green box) and MHCII negative (red box). The number of ii) MHCII negative macrophages, ii) MHCII positive macrophages and iii) alveolar macrophages, WT *n*=29, ACKR2-/- *n*=28. Statistical analysis was carried out between WT and ACKR2-/- mice. Significantly different results are indicated (*, *p* < 0.05). Error bars represent SEM.

In the lung, there are resident alveolar macrophages, and diverse populations of recruited interstitial macrophages ^28^. Here, we demonstrate that in the absence of ACKR2, there was again no difference in resident macrophages, but there were reduced numbers of the monocyte derived MHCII negative interstitial macrophage population (Figure 1c, gated as shown in Supplementary Figure 4). No differences were observed in macrophages from the pleural cavity, spleen, omentum, skin, or bone marrow (Supplementary Figure 5). Importantly, in the absence of ACKR2, the number of monocytes were unchanged in peritoneal fluid but modestly reduced in the peritoneal cavity wall. Monocyte numbers remained unchanged in all other tissues investigated (Supplementary Figure 6).

### Peritoneal immune response is impaired in ACKR2-/- mice during *Escherichia coli* infection

Next, we investigated whether reduced macrophage numbers as a result of ACKR2 deficiency had functional consequences during peritoneal infection with *E.coli*. Mice were injected intraperitoneally (i.p) with 5 x 10^6^ colony forming units (CFU) of the uropathogenic isolate CFT2073. Weight loss was comparable between WT and ACKR2-/- animals 18h after infection (Figure 2a). However, the number of bacteria was increased in the peritoneal fluid and spleen of ACKR2-/- mice indicating that bacterial elimination was impaired (Figure 2bi-ii). Bacterial burden was unchanged in the peritoneal cavity wall and liver during infection (Figure 2biiii-iv). In the absence of ACKR2, there was increased recruitment of neutrophils and monocytes to the peritoneal cavity during infection, consistent with the increased bacterial burden observed (Figure 2ci-ii). The formation of mesothelia-bound multilayered cellular aggregates is an important component of bacterial elimination during peritoneal infection ^29^. Here, we see the characteristic macrophage disappearance/ adherence reaction 18h after infection in WT and ACKR2-/- mice ^26, 29^, with a reduction in both SPMs and LPMs in the peritoneal cavity (Figure 2ciii-iv). However, in ACKR2-/- mice, increased numbers of SPMs and LPMs remain within the peritoneal cavity during infection, compared with WT, suggesting that the adherence reaction is reduced (Figure 2ciii-iv). In the peritoneal wall there is no change in SPM number between WT and ACKR2-/- mice during infection. However there are reduced numbers of LPMs in the wall of ACKR2-/- mice during infection, suggesting that the adherence reaction has been diminished, which may contribute to insufficient bacterial clearance (Figure 2dii).

**Figure 2:**
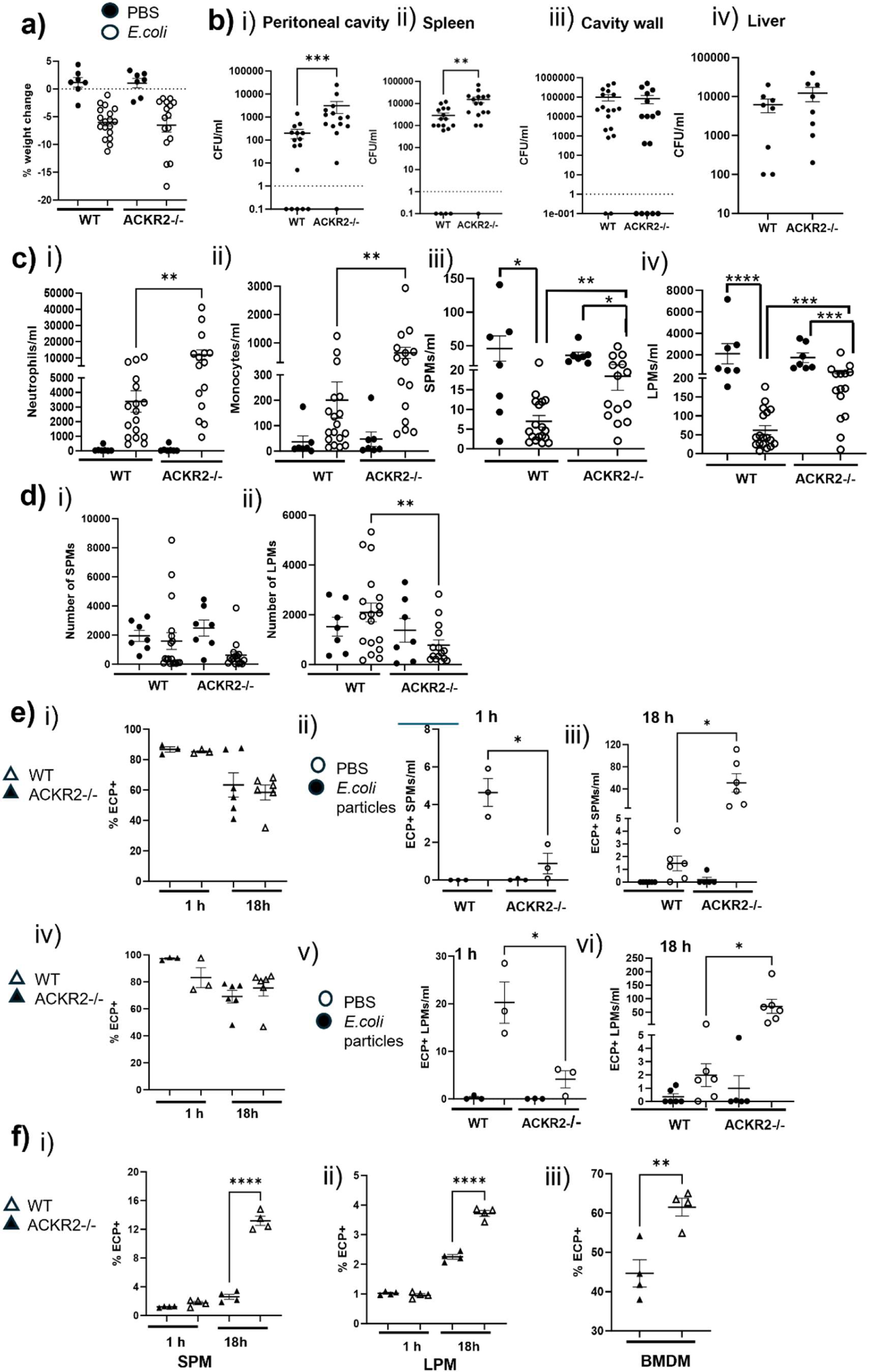
Bacterial elimination during *E.coli* infection is impaired in ACKR2-/- mice. **a)** Weight change after i.p challenge with either PBS (denoted by black circles) or 5 × 10^6^ CFU *E*. *coli* strain CFT073 (white circles), for 18 h (PBS, *n*=7 per group, WT, *n* = 18, ACKR2-/-, *n*=17). **b)** Colony forming units (CFU) were counted from **i)** peritoneal fluid, **ii)** spleen, **iii)** cavity wall, and **iv)** liver. **c)** Flow cytometry was used to determine the number of **i)** neutrophils, **ii)** monocytes, **iii)** SPMs and **iv)** LPMs in the peritoneal cavity during i.p infection. **d)** The number of **i)** SPMs and **ii)** LPMs in the peritoneal cavity wall. **e)** Percentage of uptake by **i)** SPMs and **iv)** LPMs, after i.p challenge with either PBS (denoted by black triangles) or 200 µg E.coli particles (white triangles). Number of ECP+ SPMs after **ii)** 1 h and **iii)** 18 h, and number of ECP+ LPMs after **v)** 1 h and **vi)** 18 h (PBS (black circles), *n*=3 per group, *E.coli* particles (white circles) WT 1 h, n = 3, ACKR2-/- 1 h, *n*=3, WT, ACKR2-/-, 18 h, n = 6). **f)** Percentage of uptake by **i)** SPMs, **ii)** LPMs, and **iii)** bone marrow derived macrophages (BMDM) *in vitro*, after 1 h and 18h incubation with PBS (black triangles) or 1 µg/ml E.coli particles (white triangles). Significantly different results are indicated (*, *p* < 0.05). Error bars represent SEM.

Next, we looked at the ability of macrophages to phagocytose pHRODO *E.coli* particles (ECP), which detect acidification after uptake. When particles are destroyed within the cell, fluorescent signal is no longer detected. Initially, we challenged WT and ACKR2-/- mice i.p with 200 µg of ECP. We found no difference in the percentage of peritoneal macrophages which phagocytosed particles 1h and 18h post challenge, indicating there was no defect in initial binding or uptake *in vivo* between WT and ACKR2-/- mice (Figure 2ei, iv). When we investigated the total number of macrophages which had phagocytosed particles, we found fewer ECP+ macrophages after 1 h in ACKR2-/- mice (Figure 2eii, v), likely due to decreased levels of SPMs in the peritoneal cavity, prior to infection (Figure 1aii). Later, we detected increased numbers of ECP+ macrophages in ACKR2-/- mice, suggesting that 18h after challenge, destruction of the particles is impeded (Figure 2eiii, vi). To study particle uptake without the macrophage disappearance/adherence reaction, we also carried out ECP challenge of all cells within the peritoneal lavage *in vitro*. We observed no change in the percentage of ECP+ macrophages after 1 h, but an increased proportion of ECP+ macrophages after 18h (Figure 2fi-ii), further suggesting that while binding and initial uptake are unaffected, breakdown of pHRODO particles is reduced in the macrophages from ACKR2-/- mice (Figure 2e). We also generated bone marrow derived macrophages (BMDM) from WT and ACKR2-/- mice and challenged with ECP *in vitro*. There was an increased percentage of ECP+ macrophages in BMDMs from ACKR2-/- mice, revealing the destruction of particles is again impaired (Figure 2fiii).

### ACKR2 has no detectable effect on chemokine levels in the peritoneal cavity

Next, we investigated the expression of ACKR2 by cells in the peritoneal cavity. ACKR2 expression has previously been demonstrated on peritoneal B1 cells ^30^. As ACKR2 antibodies are known to be unreliable, we employed our well validated, functional chemokine uptake assay ^17, 31, 32^. Uptake of AF647 labelled ligand, CCL22, by peritoneal lavage cells, was compared between WT and ACKR2-/- mice (Figure 3a). We confirmed that ACKR2 is expressed by B1 cell subsets, B1a, b, c and d, in the peritoneal cavity using *in vitro* and *in vivo* CCL22 uptake assays (Figure 3b). ACKR2 expression by macrophages in the peritoneal cavity was not detected using the CCL22 uptake assay (data not shown). ACKR2 expression in the peritoneal cavity wall was detected using RNAscope *in situ* hybridization. ACKR2 expression was co-localized with Lyve-1+ stromal cells, indicating that ACKR2 is expressed by LECs in the cavity wall (Fig 3c). Expression of ACKR2 by LECs in other tissues has been extensively characterized ^18, 33, 34^.

**Figure 3:**
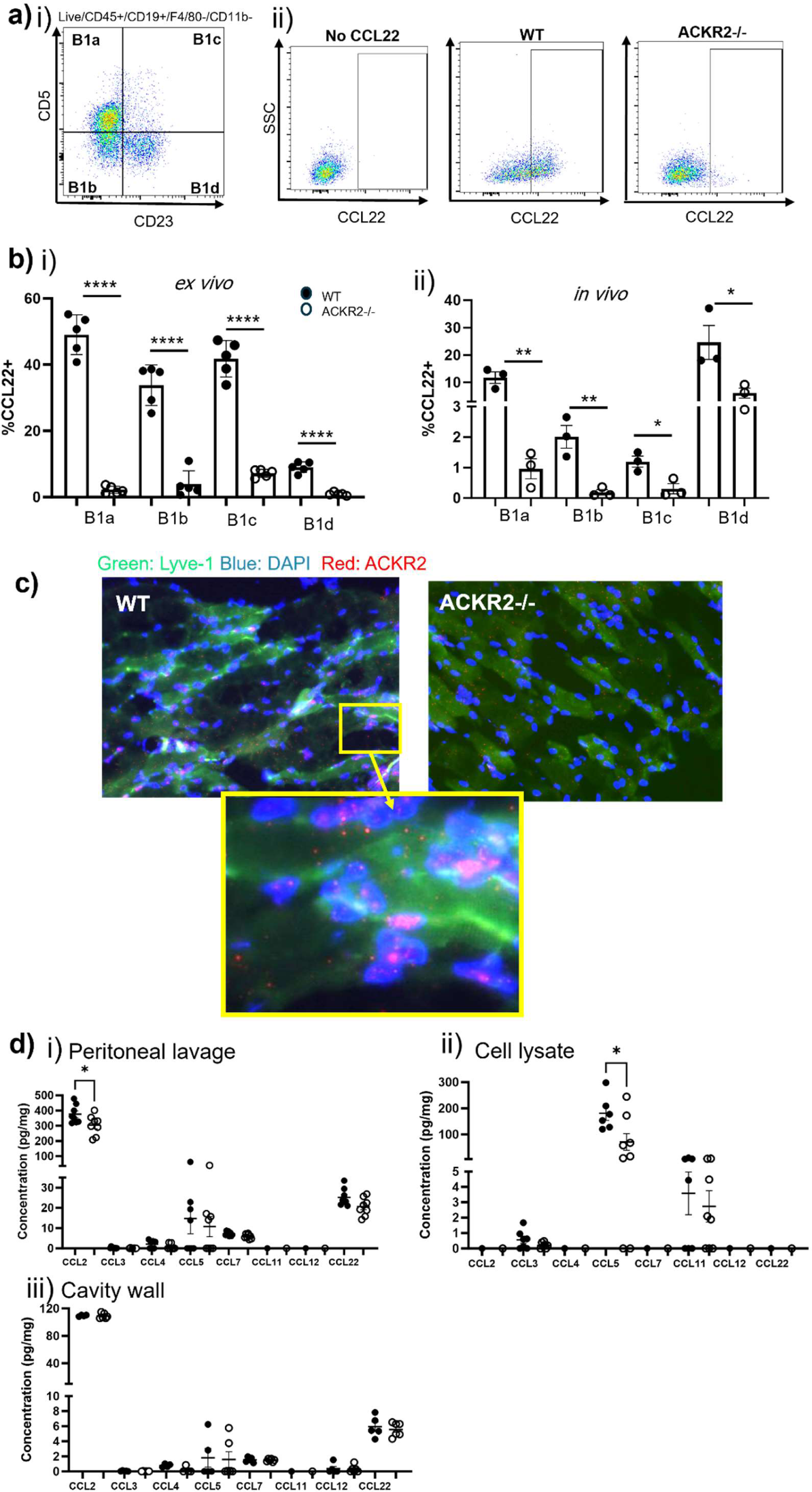
ACKR2 is expressed but has no detectable effect on chemokine scavenging in the peritoneal cavity. **a)** Flow cytometry gating to define **i)** B1 cell subsets within the peritoneal cavity based on CD5 and CD23 expression, and **ii)** ACKR2 expression by CCL22 uptake. **b)** Percentage of WT and ACKR2-/- B1 cells which are CCL22 positive after **i)** incubation with 0.25 µg/ml AF647 labelled CCL22 *in vitro* and **ii)** injection with 0.5 µg AF647 CCL22 *in vivo*. **c)** RNAscope *in situ* hybridisation of ACKR2 (red) and antibody staining of Lyve-1 in the peritoneal cavity wall. **d)** Luminex analysis was carried out to determine protein levels in **i)** peritoneal lavage fluid (*n*=8, per group), **ii)** cell lysates (WT *n*=7, ACKR2-/- *n*=8) and **iii)** cavity wall lysates (WT *n*=5, ACKR2-/- *n*=6). Significantly different results are indicated (*, *p* < 0.05). Error bars represent SEM.

Next, we looked at the effect of ACKR2 deficiency on ACKR2 ligands. We carried out Luminex protein analysis of supernatants from peritoneal lavage, and lysates from total peritoneal cells and the peritoneal cavity wall. Surprisingly, in the absence of ACKR2 we were unable to detect any defect in scavenging of any of the ACKR2 ligands investigated (Figure 3d). We found a decrease in CCL2 in the peritoneal lavage of ACKR2-/- mice, likely caused by decreased numbers of SPMs in the peritoneal cavity (Figure 1aii). Similarly, 18h after challenge with live *E.coli* or ECP, there are notable changes in ACKR2 ligands such as CCL3, CCL5, CCL12 and CCL22 but there was no defect observed in scavenging by ACKR2 (Supplementary Figure 7). These data imply that the SPM defect in ACKR2-/- mice relates to mechanisms operating outside the peritoneal cavity.

### Monocyte differentiation into macrophages is reduced in ACKR2-/- mice

As we were unable to detect local effects of ACKR2 scavenging in the peritoneal cavity, or a reduction in monocyte numbers, we next investigated whether the reduced number of macrophages in the naïve ACKR2-/- peritoneal cavity was due to altered differentiation from monocytes. We investigated the ability of bone marrow monocytes from WT and ACKR2-/- mice to differentiate into macrophages *in vivo*, by adoptively transferring either CD45.1 or CD45.2 monocytes, and tracking differentiation (Figure 4ai). To assess the ability of monocytes from WT mice to differentiate into SPMs in an ACKR2 deficient environment, we injected WT (CD45.1) monocytes i.p into either WT (CD45.2) or ACKR2-/- (CD45.2) mice. Monocytes isolated from WT mice were fully able to differentiate in the ACKR2-/- peritoneal cavity (Figure 4aii). Next, we looked at the ability of monocytes from WT (CD45.2) or ACKR2-/- (CD45.2) mice to differentiate into SPMs in a WT (CD45.1) environment. We found that monocytes from ACKR2-/- mice were less able to differentiate into SPMs, irrespective of being within a WT environment (Figure 4aiii). Again, these data point to a monocyte/macrophage-autonomous dysfunction, rather than a peritoneal dysfunction, underlying the observed phenotype.

**Figure 4:**
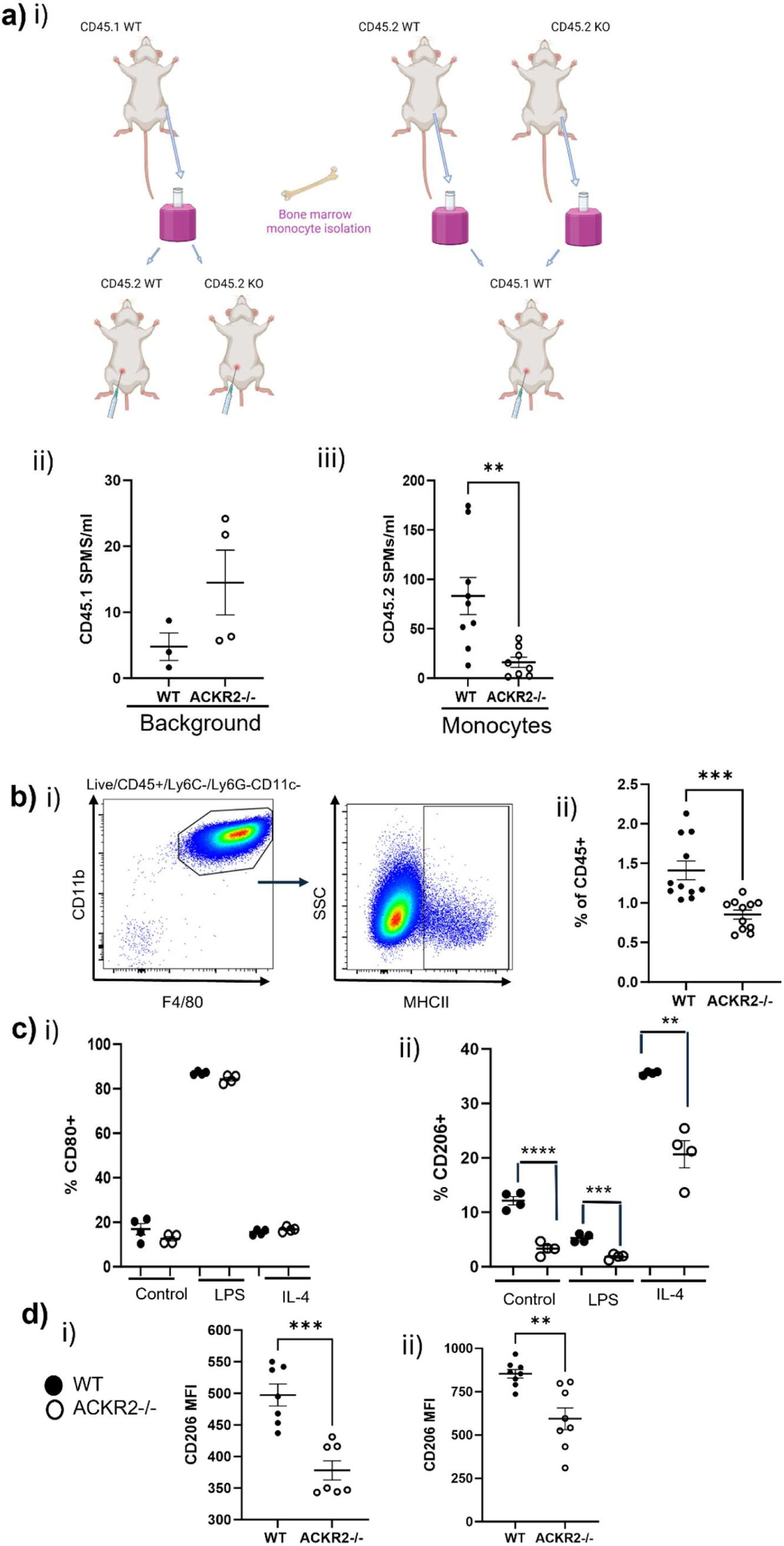
Monocyte differentiation into macrophages is reduced in the absence of ACKR2. **a) i)** Schematic of bone marrow monocyte adoptive transfer experiments. Created in BioRender.com. **ii)** 1 x 10^5^ WT CD45.1 monocytes were injected i.p into WT and ACKR2-/- CD45.2 mice (WT CD45.1 monocytes *n*=4, per group). **iii)** 1 x 10^5^ WT (*n*=9) and ACKR2-/- (*n*=8) CD45.2 monocytes were injected i.p into WT CD45.1 mice. Cells were collected 2 days later by peritoneal lavage. **b) i)** Bone marrow derived macrophage (BMDM) gating. **ii)** Percentage of MHCII expressing BMDMs (*n*=11 per group, where each point represents a culture dish, combined from at least 4 mice). **c) i)** CD80 expression and **ii)** CD206 in response to LPS and IL-4, *n*=4 per group). **d)** CD206 expression (MFI) on naïve **i)** SPMs and **ii)** MHCII negative interstitial macrophages. Significantly different results are indicated (*, *p* < 0.05). Error bars represent SEM.

We next studied the ability of monocytes from WT and ACKR2-/- bone marrow to differentiate *in vitro*. In unstimulated BMDMs generated from ACKR2-/- mice, we observed a decreased proportion which express MHCII compared with WT, after 6 d in culture (Figure 4bii). Upon stimulation, BMDMs from WT and ACKR2-/- mice had equivalent levels of CD80 expression, the classical activation marker, in response to LPS (Figure 4ci). However, we observed diminished CD206 expression by macrophages generated from ACKR2-/- bone marrow, at rest or in response to LPS or IL-4 (Figure 4 cii). Notably, we also observed lower CD206 expression in the key populations in naïve adult ACKR2-/- mice i.e. SPMs and MHCII negative interstitial macrophages (Figure 4d).

### Monocytes and macrophages are transcriptionally distinct in the absence of ACKR2

RNA sequencing of FACS sorted monocytes from the bone marrow, gated as described in Supplementary Figure 8, was carried out to investigate whether ACKR2 deficiency leads to fundamental transcriptional differences in monocyte development. Transcriptional analysis revealed substantial changes in monocytes from ACKR2-/- mice, with 113 genes upregulated and 39 downregulated (Figure 5a). Pathway analysis revealed a number of changes associated with the NFKB1 pathway, including reduced TGFB1, which is known to play an important role in alternative macrophage activation (Figure 5aiii) ^35, 36^. There are also key changes in genes, such as ALOX15, associated with the upstream regulator, signal transducer and activator of transcription 6 (STAT6) (Figure 5aiv), which is critical for IL-4 signalling ^37^. We also carried out RNA sequencing of FACS sorted SPMs from the peritoneal cavity of WT and ACKR2-/- mice. In mature macrophages, ACKR2 deficiency led to limited but distinct transcriptional changes. 16 upregulated and 15 downregulated genes were identified in SPMs from ACKR2-/- mice (Figure 5bi). Pathway analysis revealed no significantly altered pathways (Figure 5biii).

**Figure 5:**
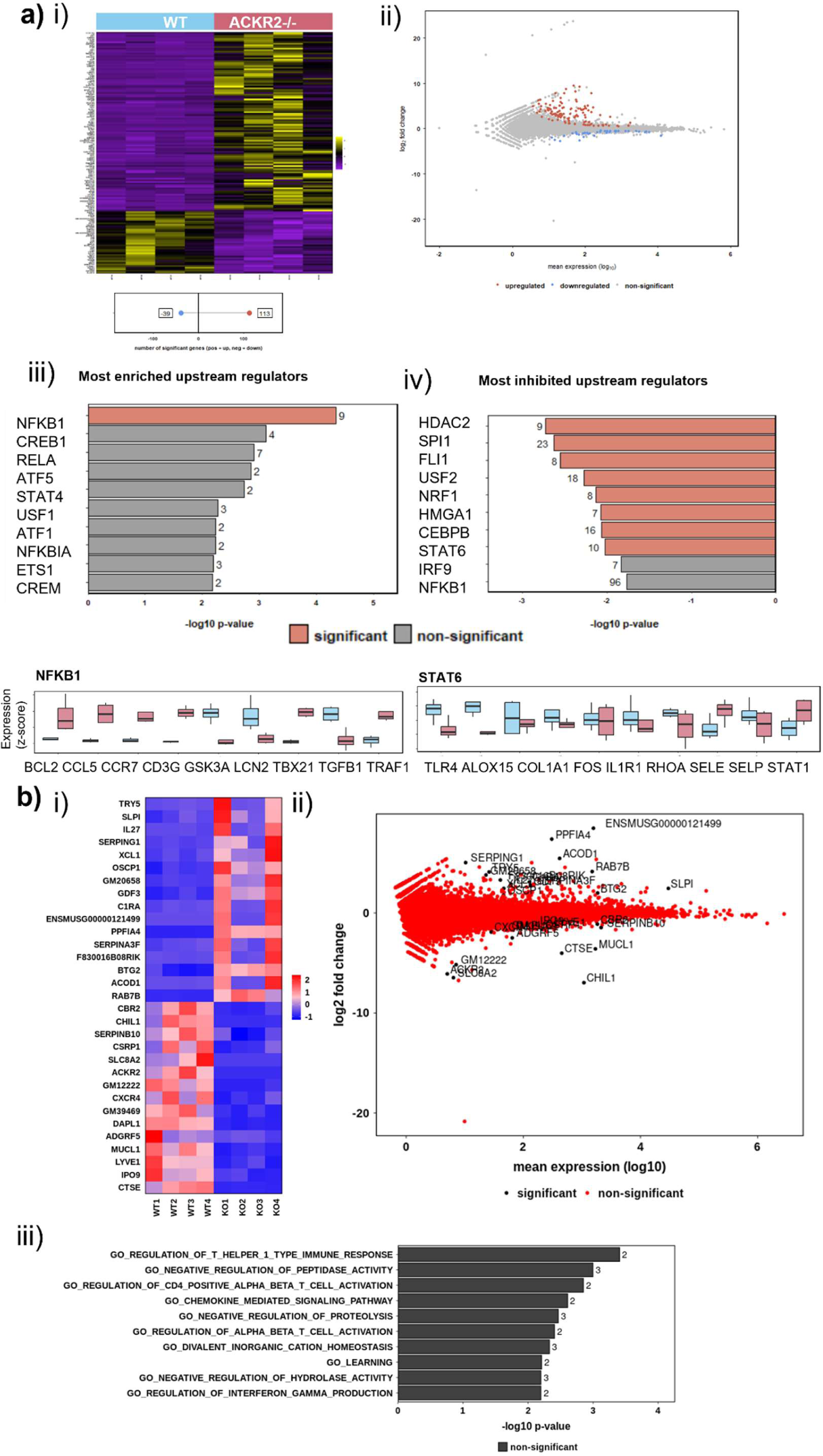
Monocytes and macrophages are transcriptionally distinct in the absence of ACKR2. **a)** Bulk RNA sequencing was carried out of FACS sorted monocytes from the bone marrow of resting WT and ACKR2-/- mice (*n*=4 per group). **i)** Hierarchically clustered heatmaps of the significantly differentially expressed genes between samples. **ii)** MA plot to show the expression of each gene and fold change between samples. Pathway analysis of **iii**) enriched upstream regulators and **iv)** inhibited upstream regulators **b)** RNA sequencing of FACS sorted SPMs from the peritoneal cavity of naïve WT and ACKR2-/- mice (*n*=4 per group). **i)** Hierarchically clustered heatmaps, **ii)** MA plot and **iii**) pathway analysis. Bioinformatic and statistical analysis was carried out using Searchlight 2 (v2.0.3), (*p*.adj < 0.05, absolute log2 fold > 0.5) ^38^.

### ACKR2 scavenges CCL11 to regulate eosinophils in the bone marrow and blood

Next, we investigated expression of ACKR2 in the bone marrow, using the AF647- CCL22 uptake assay. CCL22 uptake was increased in WT, compared with ACKR2-/- CD11b+F4/80+ macrophages, and B1b cells, indicating ACKR2 expression by these cell types in the bone marrow (Figure 6a). A previous study showed transcription of ACKR2 by HSC progenitors in the bone marrow ^14^. However, our functional uptake assay does not show ACKR2 expression by LSK or CMP cells (Supplementary Figure 9). Luminex analysis of bone marrow cell lysates and plasma from WT and ACKR2-/- mice, revealed increased levels of the ACKR2 ligand, and eosinophil chemoattractant, CCL11, suggesting that ACKR2 actively scavenges this chemokine in bone marrow and/or blood (Figure 6bi-ii). Flow cytometry revealed that in the absence of ACKR2 scavenging, the number of eosinophils was reduced in the bone marrow and increased in the blood (Figure 6ci-ii). This is consistent with deficient scavenging of CCL11 in ACKR2-/- mice, leading to increased egress of eosinophils, from the bone marrow to the blood. It has been established previously that CCL11 mobilises eosinophils within the bone marrow to enter the blood ^39^. We also demonstrated that increased CCL11 in the bloodstream leads to eosinophil egress from the bone marrow, by injecting 1 µg CCL11 intravenously (i.v) into WT mice, and measuring levels of eosinophils in the blood and bone marrow after 2 days (Figure 6d). Notably, during *E.coli* infection, the number of eosinophils within the bone marrow increased in WT mice, suggesting an important role for eosinophils in anti- bacterial immunity. In the absence of ACKR2, eosinophils do not accumulate in the bone marrow during infection, likely due to increased eosinophil egress (Figure 6e).

**Figure 6:**
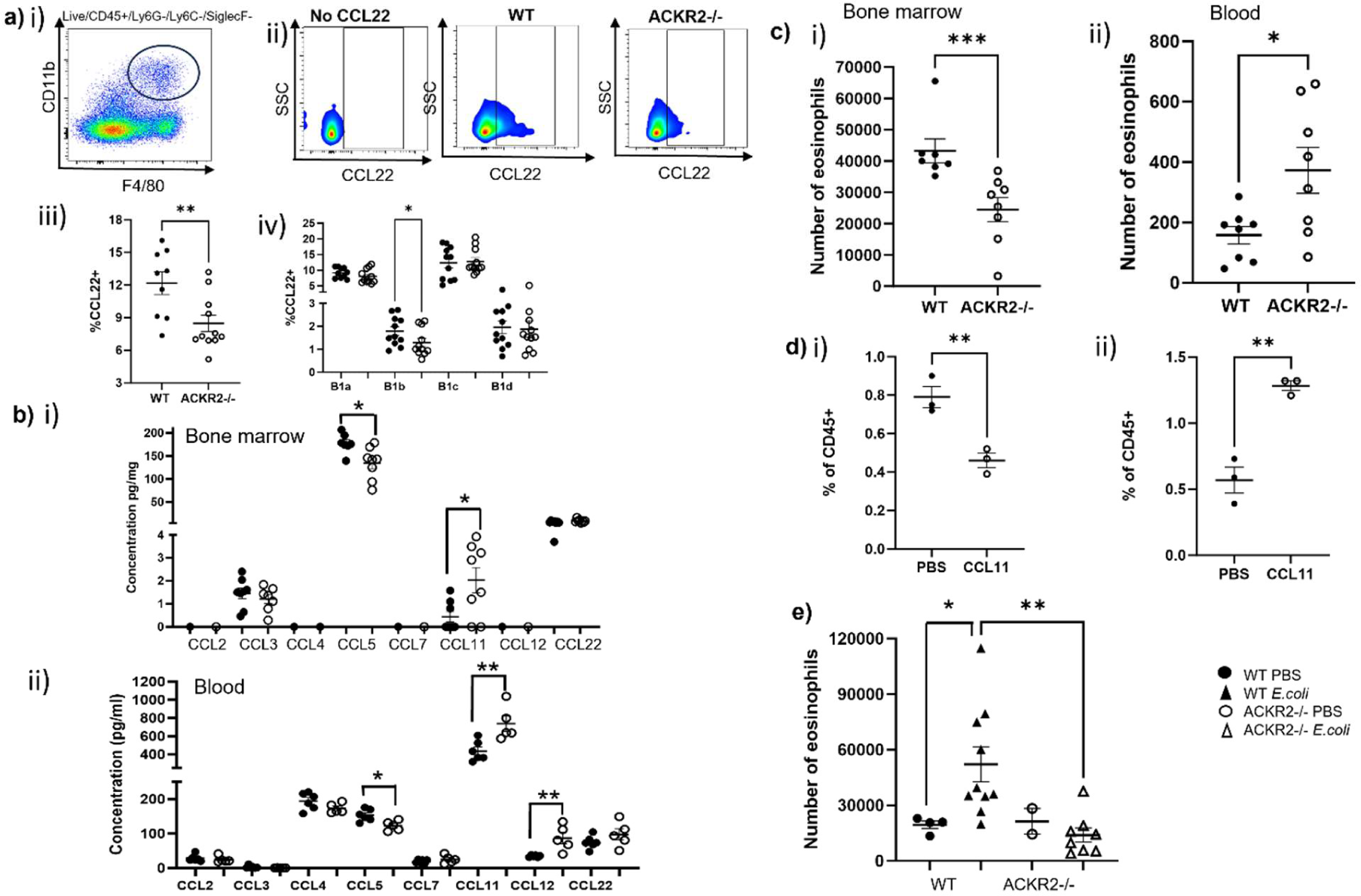
ACKR2 scavenges CCL11 to regulate levels of eosinophils in the bone marrow and blood. **a)** Flow cytometry gating strategy to define **i)** CD11b+F4/80+ macrophages within the bone marrow (BM) and **ii)** ACKR2 expression by CCL22 uptake in BM **iii)** macrophages (WT *n*=9, ACKR2-/-, *n*=11), and **iv)** B1b cells (*n*=11, per group). **b)** Luminex analysis was carried out to determine protein levels in **i)** BM cell lysates (*n*=8 per group) and **ii)** plasma (WT, *n*=5, ACKR2-/-, *n*=6). **c)** Flow cytometry was used to determine the number of eosinophils in the **i)** bone marrow and **ii)** blood of WT and ACKR2-/- mice. **d)** PBS or 1 µg of CCL11 was injected intravenously into WT mice and eosinophils were measured as a percentage of CD45+ cells in the **i)** bone marrow and **ii)** blood. **e)** The number of eosinophils in the bone marrow 18h after infection, in WT and ACKR2-/- mice was measured by flow cytometry. Significantly different results are indicated (*, *p* < 0.05). Error bars represent SEM.

### Eosinophils promote monocyte to macrophage differentiation

We have shown that eosinophil retention in the bone marrow is ACKR2 dependent, via scavenging of CCL11 (Figure 6). Therefore, we next investigated whether eosinophils and monocytes interact within the bone marrow. Firstly, we used iREP mice, a reporter mouse strain we have recently generated, where each of the inflammatory chemokine receptors CCR1, 2, 3 and 5 are discretely labelled ^23^. We were able to visualise CCR2+ monocytes and CCR3+ eosinophils in close proximity within the bone marrow (Figure 7a). To investigate if eosinophils promote monocyte to macrophage differentiation, we added FACS sorted WT eosinophils on d 0, to WT and ACKR2-/- bone marrow cultures *in vitro*. When stimulated with IL-4, eosinophil treated BMDMs from ACKR2-/- mice were able to respond to IL-4 at equivalent levels to WT (Figure 7b). In addition, treatment with eosinophils restored the ability of BMDMs from ACKR2-/- mice to destroy ECP after phagocytosis (Figure 7c).

**Figure 7:**
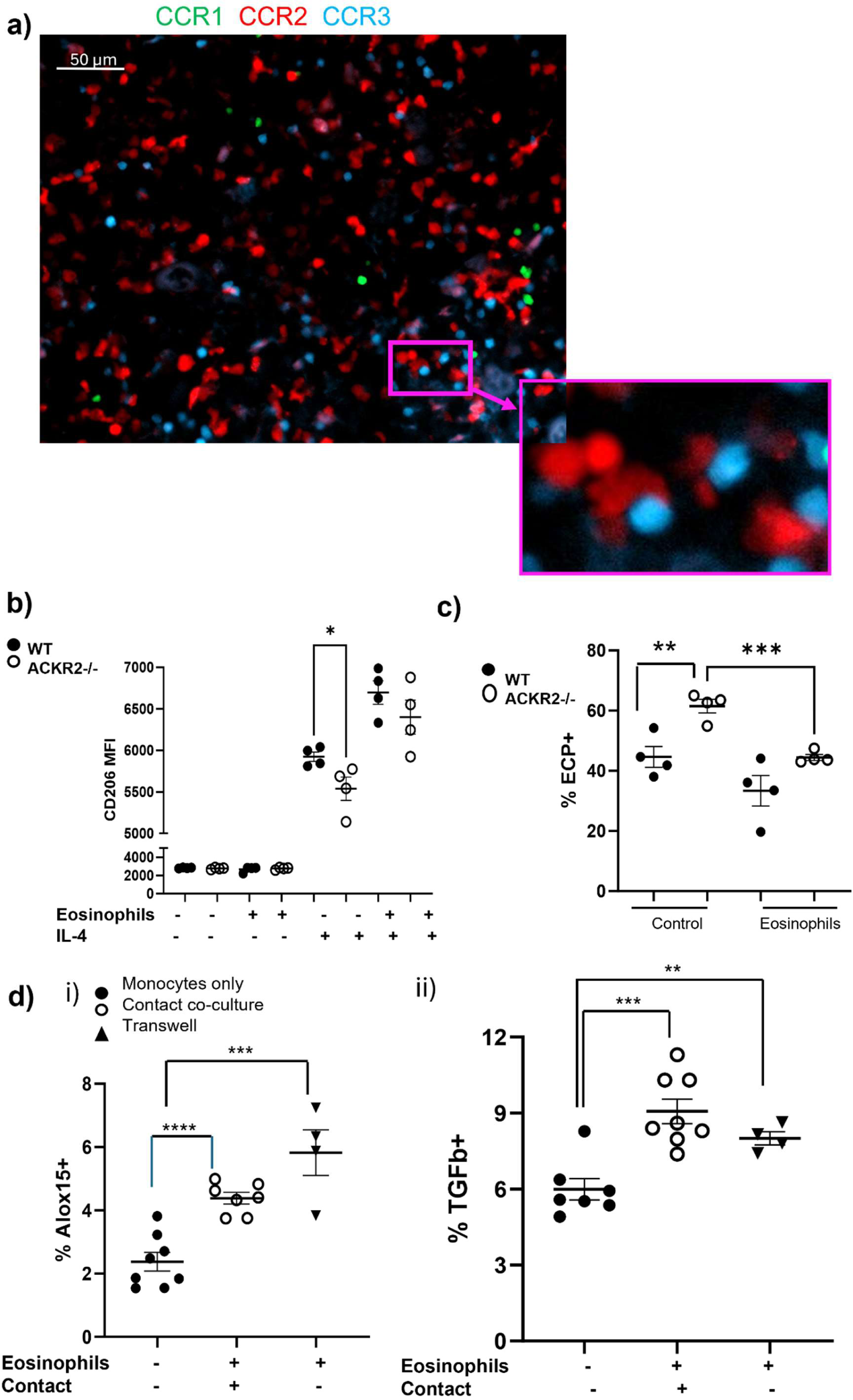
Eosinophils promote monocyte to macrophage differentiation. **a)** Bone marrow from iREP mice was imaged at 20 x magnification. CCR1 expression is green, CCR2 is red, and CCR3 is blue. **b)** WT (black circles) and ACKR2-/- (white circles) BMDMs were generated with and without FACS sorted WT eosinophils on day 0, and with and without IL-4 treatment for 18h on day 5 (*n*=4 per group). **c)** Percentage of pHRODO *E.coli* particle positive BMDMs with and without eosinophil treatment (*n*=4 per group). **d)** Intracellular **i)** ALOX15 and **ii)** TGF-β production after 18h co-culture of FACS sorted monocytes and eosinophils (no eosinophils, *n*=7, eosinophils and monocytes with contact, *n*=8, without contact, *n*=4). Significantly different results are indicated (*, *p* < 0.05). Error bars represent SEM.

We have identified key transcriptional changes in monocytes from the bone marrow of ACKR2-/- mice, where monocytes had less exposure to eosinophils, such as reduced transcription of TGFB1 and ALOX15 (Figure 5aiii-iv). To investigate whether increased exposure to eosinophils was able to drive expression of these genes in monocytes, we co-cultured FACS sorted monocytes and eosinophils for 18h *in vitro*. Upon co-culture with eosinophils, monocytes produced more TGFB1 and ALOX15, detected by intracellular flow cytometry (Figure 7d). This process was not contact dependent, as eosinophils cultured in a transwell system were also able to drive the gene expression changes in monocytes (Figure 7d).

## DISCUSSION

Monocyte derived macrophages are a key component of the immune response, and are found in most tissues throughout the body ^3^. Monocytes are recruited from the bone marrow and differentiate into macrophages at distal sites. Here we describe, a previously unknown role for eosinophils, classically known as mediators of type 2 immunity, in educating monocytes within the bone marrow, prior to recruitment into tissues.

We demonstrate a novel interaction between eosinophils and monocytes in the bone marrow, indirectly controlled by the atypical chemokine receptor ACKR2. ACKR2 regulates eosinophils within the bone marrow by scavenging CCL11, a ligand shared with the dominant eosinophil chemokine receptor CCR3 ^22, 23^. CCL11 is a potent chemoattractant for eosinophils, and its role in the mobilisation of eosinophils from the bone marrow to the blood, has long been known ^39^. In the absence of ACKR2, CCL11 is not scavenged in the bone marrow or the blood, and increased eosinophil egress is observed. As a result of this, eosinophils and monocytes interact less within the bone marrow, leading to changes in monocyte gene expression. Monocytes from ACKR2-/- mice are able to travel to the tissues in equivalent numbers, but their ability to differentiate into macrophages is fundamentally altered, in the lung, peritoneal cavity and peritoneal wall. This mechanism has been illustrated as a schematic in Figure 8. In this study, we did not observe changes in any of the other tissues we investigated, such as the skin, omentum, or the spleen. Surprisingly, we also did not observe any reduction in pleural cavity SPMs, which are similar to peritoneal SPMs ^27^, suggesting that these cells have an altered differentiation pathway. Importantly, BMDMs generated from ACKR2-/- mice exhibit the key differences observed in tissue specific macrophages, such as reduced MHCII expression, impaired breakdown of pHRODO *E.coli* particles, and diminished CD206 expression. This suggests that the effect on monocyte derived macrophages observed in ACKR2-/- mice at rest is stable *ex vivo* and may not be limited to the peritoneal cavity and lung.

**Figure 8:**
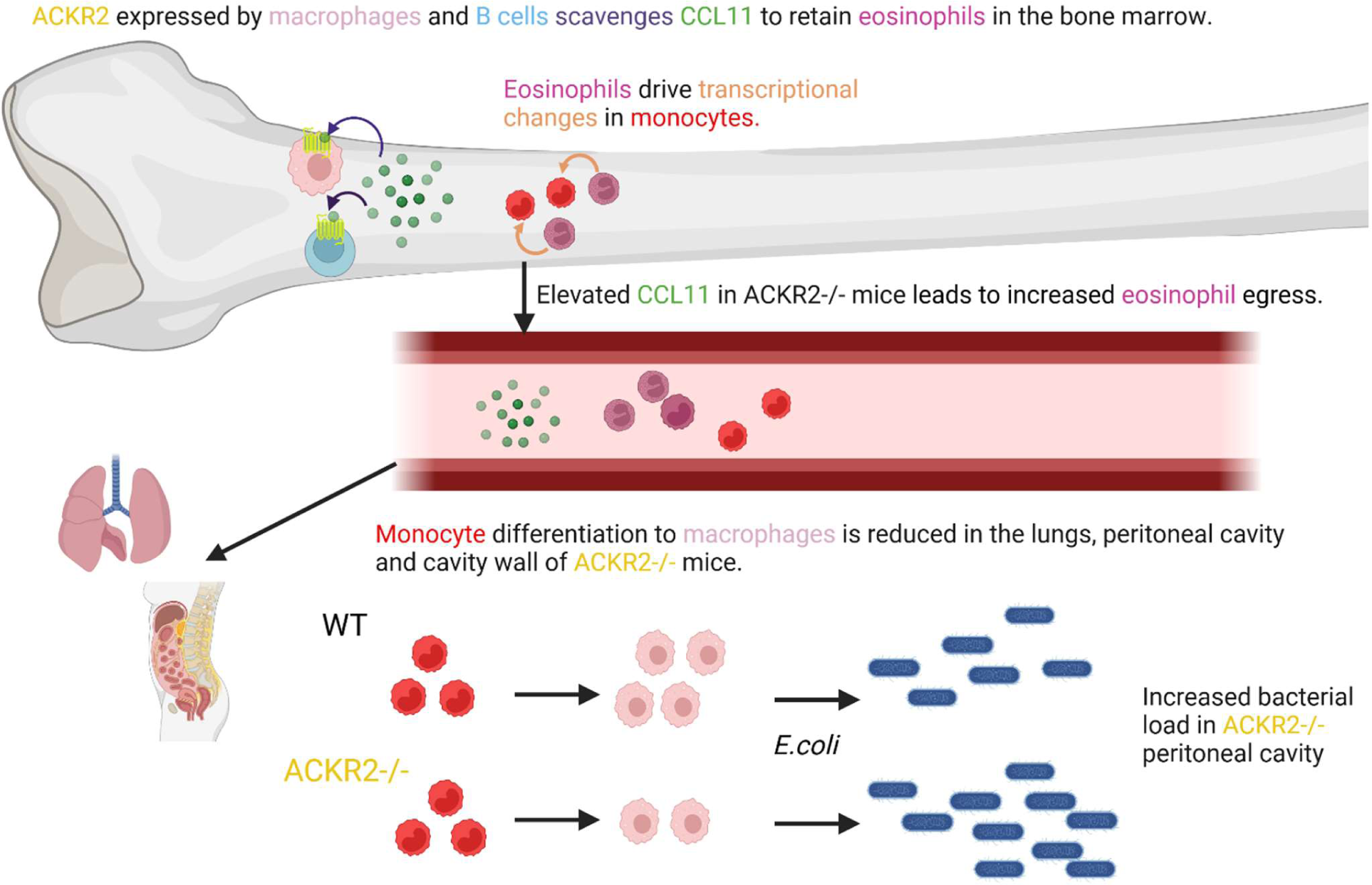
Schematic to illustrate how ACKR2 promotes monocyte to macrophage differentiation by limiting eosinophil egress from the bone marrow. Created in BioRender.

Reduced differentiation of monocytes to macrophages in the peritoneal cavity of ACKR2-/- mice has clear functional implications, exemplified by increased bacterial load in the peritoneal cavity during infection. Our transcriptional analysis of SPMs revealed highly specific gene changes, which may contribute to the reduced functionality of macrophages from ACKR2-/- mice during peritoneal infection. SPMs from ACKR2-/- mice have reduced transcription of CTSE and Chil1 (Figure 5bi). CTSE-/- macrophages have decreased bacteriocidal activity, and CTSE-/- mice are highly susceptible to infection with *Staphylococcus aureus* and *Porphyromonas gingivalis* ^40, 41^. Similarly, Chil1-/- mice are more susceptible to *Pseudomonas aeruginosa* infection ^42^. Additionally, the formation of cellular aggregates of resident macrophages and mesothelia within the peritoneum is important for bacterial killing^29^. Here, we observed reduced numbers of resident macrophages in the wall of ACKR2-/- mice during infection, which may contribute to insufficient bacterial elimination. ACKR2 has previously been shown to be important during infection with *Mycobaterium tuberculosis,* attributed to increased chemokine and cytokine production in ACKR2-/- mice ^43^. Here we reveal a lung macrophage defect in ACKR2-/- mice, which may have contributed to increased pathology. In a caecal ligation and puncture (CLP) sepsis model there is also evidence of a protective role of ACKR2 ^44^.

Previously, expression of ACKR2 by osteoclasts and osteoblasts has been demonstrated during bone remodeling ^45^. Here we show for the first time, ACKR2 expression by macrophages and B1b cells in the bone marrow, at rest. Previously, ACKR2 expression has mainly been identified in stromal populations, such as LECs and fibroblasts, throughout the body, or by peritoneal B1 cells ^9^. Notably, several human GWAS studies have revealed SNPs in the ACKR2 gene, which are associated with monocyte (rs2228467, rs1366045, rs136046, rs219224, and rs2228468) and eosinophil counts (rs2228467, rs7433284, rs9872570), lending further support to the idea that ACKR2 plays an important role in controlling human monocyte and eosinophil interactions ^46, 47, 48, 49, 50^.

Several studies have revealed a role for eosinophils in polarizing macrophages towards an M2 or alternatively activated phenotype ^51, 52, 53^. The transfer of eosinophils during collagen induced arthritis increased the proportion of CD163+ macrophages and TGF-β production in arthritic joints ^51^. In adipose tissue, alternative macrophage activation was impaired in the absence of eosinophils ^53^. Here, we show that eosinophils and monocytes are located in close proximity within the bone marrow, and eosinophils transcriptionally alter monocytes, which is necessary for full CD206 expression in response to stimulation with IL-4. We also provide a novel ACKR2-dependent control mechanism through CCL11 scavenging. Notably, co-culture of eosinophils and monocytes was sufficient to recapitulate transcriptional changes observed in ACKR2-/- monocytes. We have shown that this process does not require direct contact between the cell types, which suggests that secreted factors or degranulation products may mediate the effect. Future work will be important to determine the nature of the factor(s) responsible. During bacterial infection, we observed an increase in eosinophils within the WT but not ACKR2-/- bone marrow. Taken together with the increased bacterial load in ACKR2-/- mice, eosinophil/monocyte interactions appear to represent a key component of the anti- bacterial immune response.

Thus, ACKR2 is a key regulator of eosinophil-driven monocyte education in the bone marrow which is required for full monocyte differentiation and macrophage function within the tissues.

## Materials and Methods

### Study design

The objective of this study was to reveal how ACKR2 regulated the numbers of macrophages in different tissues, and the response to bacterial infection. A combination of *in vitro* and *in vivo* approaches have been used. All mouse experiments were carried out using age and sex matched animals. Unless otherwise stated each point represents an individual mouse. The number of replicates, *n* has been reported for each experiment. At least three biological replicates per experimental group were used. Required sample size was calculated based on previous similar experiments.

### Animals

Animal experiments were carried out under the auspices of a UK Home Office Project Licence. All experiments conformed to animal care and welfare protocols, approved by the local ethical review committee at the University of Glasgow. Wild-type (WT) C57BL/6J mice were purchased from Charles River Laboratories. CD45.1 ^54^, ACKR2-/- ^55^ and iREP ^23^ mice were bred in the specific pathogen-free facility of the University of Glasgow Wolfson Research Facility (WRF).

### Flow cytometry

Flow cytometry and cell sorting experiments were carried out according to established guidelines ^56^. Dead cells were excluded using Fixable Viability Dye eFluor 506 (Thermo Fisher Scientific). Cell suspensions were blocked with TruStain FcX (Biolegend) at 1:200. Antibodies were used at a dilution of 1 in 200 in FACS buffer (PBS, 1% foetal bovine serum (FBS), and 2 mM EDTA (Thermo Fisher Scientific)). Anti-mouse antibodies of the following clones were used on various fluorochromes (Biolegend), CD45 (30-F11), CD11b (M1/70), F4/80 (BM8), Siglec F (S17007L), Ly6C (HK1·4), CD11c (N418), MHCII (M5/114·15·2), and CD206 (C068C2), CD5 (53-7.3), CD23 (B3B4), CD45.1 (A20), CD45.2 (104), CD80 (2D10), Sca1 (D7), c-kit (2B8), CD34 (MEC14.7), TGF-β1 (TW7-16B4). Ly6G BUV395 (1A8) and CD19 BUV805 (1D3) were obtained from BD Biosciences. Lyve-1- Alexa Fluor 488 (223322) was from R&D Systems, and ALOX15 Alexa Fluor 488 (AA 581-662) from Antibodies online. Cell suspensions were washed with FACS buffer and fixed with 2% PFA. Intracellular staining was carried out using the Cyto Fast Fix Perm buffer set (Biolegend), as described in the manufacturer’s instructions. Flow cytometry was performed using a Fortessa or Celesta (BD Biosciences) and analysed using FlowJo V10. FACS sorting was carried out using a FACSaria III (BD Biosciences).

### Cell isolation

Peritoneal lavage was carried out by injecting 10 ml PBS containing EDTA into the peritoneal cavity of euthanized mice and collecting the fluid with a syringe. Bone marrow cells were obtained by dissecting the hind limbs, removing the muscle, cutting the femur and tibia bones and flushing with 1 ml PBS containing EDTA. Red blood cells (RBC) were lysed in 1 mL RBC lysis buffer (Invitrogen) for 1 min, and then quenched in 10 ml PBS. Blood was collected by cardiac puncture into a 1 ml syringe containing 50 µl 0.5 M EDTA. Cells were washed in FACS buffer and analysed by flow cytometry.

### Tissue Digestion

Lungs were removed and chopped finely, before digestion with 0.8 mg/mL Dispase II, 0.2 mg/mL Collagenase P, and 0.1 mg/mL DNase I (Roche) in HBSS shaken at 800 rpm, at 37°C for 45 min. Lungs were filtered through a 40 µm EASYstrainer mesh (Greiner Bio-One), and RBC were lysed. Peritoneal cavity wall was excised and chopped finely, before digestion with 1ml of Liberase at 0.44 Wunch units per sample, for 1 h at 37 °C, 1000 rpm using a Thermoshaker. Cell suspension was filtered through a 100µm EASYstrainer, RBCs were lysed and washed with 10 ml PBS.

### BMDM generation

Bone marrow cells were isolated from the femurs of at least 4 WT and ACKR2-/- mice. Cells were grown in media consisting of 433 ml GMEM (Glasgow Minimum Essential Medium), 50 ml FBS (foetal bovine serum), 20 ng/ml mouse M-CSF Recombinant Protein (Peprotech, 5 ml nonessential amino acids, 5 ml L-glutamine, 5 ml sodium pyruvate, 0.1mM β-mercaptoethanol and 1 ml of Primocin. Bone marrow was collected, unwanted debris removed using a 70 μm filter, and centrifuged at 300 g for 5 min. The supernatant was discarded, the cell pellet was resuspended in 1 ml of ACK lysis buffer and incubated at room temperature for 1 minute. Next, cells were centrifuged at 300g for 5 minutes and resuspended in 1 ml of BMDM media. Cell numbers were determined using 1:1 dilution with Trypan blue, and seeded at a density of 5 x 10^5^ in 35 mm Petri dishes with BMDM media. On day 3, media was replaced. BMDMs were stimulated on day 5 with either IL-4 (Biolegend) at 20 ng/ml, or LPS at 10ng/ml (eBioscience). In some experiments, 20,000 FACS sorted eosinophils were added on day 0 at a ratio of 1:25.

### Monocyte and Eosinophil co-culture

Monocytes and eosinophils were purified by FACS sorting using the FACSAria III (BD Biosciences), and cultured together at a ratio of 10:1, for 18h. Millicell 96 well Plate with 0.4 µm pore size (Millipore, Sigma) was used to assess whether gene expression changes were contact dependent. After 18h cells were stained and assessed by flow cytometry.

### Peritoneal bacterial infection

Mice between 7 - 9 weeks of age were injected intraperitoneally (i.p) with 5 × 10^6^ CFU of *E*. *coli* (UPEC isolate CFT073) as described previously ^57^. Bacteria were grown overnight in Luria-Bertani (LB) medium, then sub-cultured the following day and grown to log phase for injection (OD_600_ = 0·5, 5 × 10^8^ CFU/ml). Mice were monitored for weight loss and clinical signs, culled 18h later, and lavages were collected. For CFU analysis, 1 ml of the lavage fluid was centrifuged at 13000 g and resuspended in 50 µl for serial dilution. To determine CFU values from the spleen, peritoneal cavity wall and liver, tissue was disrupted using stainless steel beads and a Tissuelyser LT (Qiagen) for 10 min at a rate of 50 s^−1^. CFU were determined by plating serial dilutions on LB agar.

### Sterile inflammation of the peritoneal cavity

Mice were injected i.p with either 200 µl PBS or 200 µg of pHRODO Bioparticles in 200ul PBS (*E.coli* K-12 strain, Thermo Fisher Scientific). After 18h, mice were culled, peritoneal lavage was collected and processed for cellular analysis.

### Adoptive transfer experiments

Monocytes were purified from bone marrow cells isolated from 4 mice, by magnetic sorting using the EasySep™ Mouse Monocyte Isolation Kit (Stem Cell Technologies), as described in the manufacturer’s instructions). 1 x 10^5^ monocytes were injected i.p into mice, and cells were collected 2 days later by peritoneal lavage.

### Imaging

Bone marrow from iREP mice were visualized at 20 x magnification on an Axioimager. Samples were prepared using a tape strip methodology developed by Dr Tadafumi Kawamoto ^58^. Peritoneal wall was fixed using 4% PFA overnight, then placed into 30% sucrose overnight before being frozen in OCT. ACKR2 expression in the peritoneal wall was detected by RNAscope *in situ* hybridization using a specific probe, as described previously ^16, 17^. Co-staining with Lyve-1 Alexa Fluor 488 labelled antibody (R&D systems) was carried out at 1:200 dilution in 1% BSA in PBS containing 0.1% Tween-20 (Sigma), overnight at 4°C.

### Protein samples

Peritoneal wall tissue was frozen in liquid nitrogen, crushed using a mortar and pestle, and resuspended in dH_2_O containing protease inhibitors (Pierce). Plasma was obtained from blood samples collected in 50 µl 0.5 M EDTA, and centrifuged at 1500 g for 10 min at 4 °C. Peritoneal lavage cells were centrifuged at 400 g and supernatants were removed for analysis. Cell pellets from the peritoneal lavage and bone marrow were lysed using Lysis 2 buffer (R&D systems), containing HALT protease inhibitors (Thermo Fisher Scientific). Protein concentrations were determined using a Magnetic Luminex Multiplex assay (custom designed, R&D Systems), as described in the manufacturer’s instructions, and read using a Bio-Rad Luminex-100 machine. Data was normalised to the total protein concentration of tissue samples, as determined by a BCA assay (Pierce).

### CCL22 Uptake assay

ACKR2 expression was determined by comparing uptake of Alexa Fluor 647 (AF647) labelled CCL22 by WT and ACKR2-/- cells, as described previously ^17, 19, 32^. *In vivo* ACKR2 expression was assessed by injecting either PBS or 0.5 µg of AF647 CCL22 i.p into WT and ACKR2-/- animals. 1 h later, cells were collected by peritoneal lavage stained and analysed by flow cytometry. *In vitro* uptake assays were performed by incubating WT and ACKR2-/- cells with 0.25 µg/ml AF647 CCL22, at 37 °C and 5% CO_2_ for 1 h. Cells were then stained as usual at 4 °C and analysed by flow cytometry.

### Transcriptional Analysis

Monocytes from the bone marrow and SPMs from the peritoneal cavity were FACS sorted using a FACSAria III (BD Biosciences) directly into RLT buffer (Qiagen). For each individual sample, cells were sorted from 5 mice. RNA was extracted using an RNeasy micro kit with on column DNase treatment (Qiagen) as described in the manufacturer’s instructions. RNA sequencing was carried out by Novogene for RNA QC, mRNA library preparation (poly-A enrichment) and Illumina Sequencing (PE150) with SMARTer sequence amplification. Bioinformatic analysis was carried out using Searchlight 2 (v2.0.3) ^38^.

### Statistical analysis

Data were analysed using GraphPad Prism version 10.1.2. Normality was assessed using Shapiro–Wilk and Kolmogorov–Smirnov tests. Normally distributed data was analysed using two-tailed, unpaired t-tests. Where data was not normally distributed, Mann–Whitney tests were used. Significance was defined as *p* < 0·05, *. Error bars indicate standard error of the mean (SEM).

## Acknowledgements

We thank the University of Glasgow’s animal facility staff for the care of our animals and Cellular Analysis Facility staff Diane Vaughn and Alana Hamilton for technical assistance. We thank Zhenyu Li and Mariane Jeong for technical assistance. We also thank Madeleine Cunningham and Jennifer Barrie for lab management and helpful discussions. We thank Derek Gilroy and Michelle Sugimoto for advice.

## Author Contributions

GJW conceived the study, performed experiments, analysed data and wrote the paper. LKF, ZP, EP, AF, RB, KML, LF, and HM performed experiments, reviewed and edited the paper. JJC analysed data. GJG acquired funding, conceived the study and wrote the paper.

## References

1. Mass, E., Nimmerjahn, F., Kierdorf, K. & Schlitzer, A. Tissue-specific macrophages: how they develop and choreograph tissue biology. Nature Reviews Immunology 23, 563–579 (2023).

2. Yona, S. et al. Fate Mapping Reveals Origins and Dynamics of Monocytes and Tissue Macrophages under Homeostasis. Immunity 38, 79–91 (2013).

3. Gordon, S., Plüddemann, A. & Martinez Estrada, F. Macrophage heterogeneity in tissues: phenotypic diversity and functions. Immunological Reviews 262, 36–55 (2014).

4. Rot, A. & von Andrian, U.H. Chemokines in innate and adaptive host defense: basic chemokinese grammar for immune cells. Annu Rev Immunol 22, 891–928 (2004).

5. Geissmann, F., Jung, S. & Littman, D.R. Blood Monocytes Consist of Two Principal Subsets with Distinct Migratory Properties. Immunity 19, 71–82 (2003).

6. Nibbs, R.J. & Graham, G.J. Immune regulation by atypical chemokine receptors. Nat Rev Immunol 13, 815–829 (2013).

7. Bachelerie, F. et al. International Union of Basic and Clinical Pharmacology. LXXXIX. Update on the Extended Family of Chemokine Receptors and Introducing a New Nomenclature for Atypical Chemokine Receptors. Pharmacological Reviews 66, 1–79 (2014).

8. Gordon, S. & Taylor, P.R. Monocyte and macrophage heterogeneity. Nat Rev Immunol 5, 953–964 (2005).

9. Comerford, I. & McColl, S.R. Atypical chemokine receptors in the immune system. Nature Reviews Immunology 24, 753–769 (2024).

10. Shams, K. et al. Spread of Psoriasiform Inflammation to Remote Tissues Is Restricted by the Atypical Chemokine Receptor ACKR2. Journal of Investigative Dermatology 137, 85–94 (2017).

11. Codullo, V. et al. An investigation of the inflammatory cytokine and chemokine network in systemic sclerosis. Annals of the Rheumatic Diseases 70, 1115–1121 (2011).

12. Vetrano, S. et al. The lymphatic system controls intestinal inflammation and inflammation-associated Colon Cancer through the chemokine decoy receptor D6. Gut 59, 197–206 (2010).

13. Hansell, C.A.H. et al. The Atypical Chemokine Receptor Ackr2 Constrains NK Cell Migratory Activity and Promotes Metastasis. The Journal of Immunology 201, 2510–2519 (2018).

14. Massara, M. et al. ACKR2 in hematopoietic precursors as a checkpoint of neutrophil release and anti-metastatic activity. Nature Communications 9, 676 (2018).

15. Hayes, A.J. et al. Enhanced CCR2 expression by ACKR2-deficient NK cells increases tumoricidal cell therapy efficacy. Journal of Leukocyte Biology (2024).

16. Wilson, G.J. et al. Chemokine receptors coordinately regulate macrophage dynamics and mammary gland development. Development 147 (2020).

17. Wilson, G.J. et al. Atypical chemokine receptor ACKR2 controls branching morphogenesis in the developing mammary gland. Development 144, 74–82 (2017).

18. Lee, K.M., Danuser, R., Stein, J.V., Graham, D., Nibbs, R.J.B. & Graham, G.J. The chemokine receptors ACKR2 and CCR2 reciprocally regulate lymphatic vessel density. The EMBO Journal 33, 2564–2580-2580 (2014).

19. Lee, K.M. et al. Placental chemokine compartmentalisation: A novel mammalian molecular control mechanism. PLOS Biology 17, e3000287 (2019).

20. Arnold, I.C. & Munitz, A. Spatial adaptation of eosinophils and their emerging roles in homeostasis, infection and disease. Nature Reviews Immunology (2024).

21. Cosway, E.J. et al. Eosinophils are an essential element of a type 2 immune axis that controls thymus regeneration. Science Immunology 7, eabn3286 (2022).

22. Dyer, D.P. et al. Chemokine Receptor Redundancy and Specificity Are Context Dependent. Immunity 50, 378–389.e375 (2019).

23. Medina-Ruiz, L. et al. Analysis of combinatorial chemokine receptor expression dynamics using multi-receptor reporter mice. eLife 11, e72418 (2022).

24. Cassado, A.d.A., D’Império Lima, M.R. & Bortoluci, K.R. Revisiting Mouse Peritoneal Macrophages: Heterogeneity, Development, and Function. Frontiers in Immunology 6 (2015).

25. Ghosn, E.E.B. et al. Two physically, functionally, and developmentally distinct peritoneal macrophage subsets. Proceedings of the National Academy of Sciences 107, 2568–2573 (2010).

26. Louwe, P.A. et al. Recruited macrophages that colonize the post-inflammatory peritoneal niche convert into functionally divergent resident cells. Nature Communications 12, 1770 (2021).

27. Kim, K.-W. et al. MHC II+ resident peritoneal and pleural macrophages rely on IRF4 for development from circulating monocytes. Journal of Experimental Medicine 213, 1951–1959 (2016).

28. Li, X. et al. Coordinated chemokine expression defines macrophage subsets across tissues. Nature Immunology 25, 1110–1122 (2024).

29. Vega-Pérez, A. et al. Resident macrophage-dependent immune cell scaffolds drive anti-bacterial defense in the peritoneal cavity. Immunity 54, 2578–2594.e2575 (2021).

30. Hansell, C.A. et al. Universal expression and dual function of the atypical chemokine receptor D6 on innate-like B cells in mice. Blood 117, 5413–5424 (2011).

31. Ford, L.B., Cerovic, V., Milling, S.W.F., Graham, G.J., Hansell, C.A.H. & Nibbs, R.J.B. Characterization of Conventional and Atypical Receptors for the Chemokine CCL2 on Mouse Leukocytes. The Journal of Immunology 193, 400–411 (2014).

32. Hansell, C.A.H., Love, S., Pingen, M., Wilson, G.J., MacLeod, M. & Graham, G.J. Analysis of lung stromal expression of the atypical chemokine receptor ACKR2 reveals unanticipated expression in murine blood endothelial cells. European Journal of Immunology 50, 666–675 (2020).

33. McKimmie, C.S. et al. An analysis of the function and expression of D6 on lymphatic endothelial cells. Blood 121, 3768–3777 (2013).

34. Nibbs, R.J.B. et al. The β-Chemokine Receptor D6 Is Expressed by Lymphatic Endothelium and a Subset of Vascular Tumors. The American Journal of Pathology 158, 867–877 (2001).

35. Zhang, F. et al. TGF-β induces M2-like macrophage polarization via SNAIL-mediated suppression of a pro-inflammatory phenotype. Oncotarget 7, 52294–52306 (2016).

36. Gong, D., Shi, W., Yi, S.-j., Chen, H., Groffen, J. & Heisterkamp, N. TGFβ signaling plays a critical role in promoting alternative macrophage activation. BMC Immunology 13, 31 (2012).

37. Takeda, K. et al. Essential role of Stat6 in IL-4 signalling. Nature 380, 627–630 (1996).

38. Cole, J.J. et al. Searchlight: automated bulk RNA-seq exploration and visualisation using dynamically generated R scripts. BMC Bioinformatics 22, 411 (2021).

39. Palframan, R.T., Collins, P.D., Williams, T.J. & Rankin, S.M. Eotaxin induces a rapid release of eosinophils and their progenitors from the bone marrow. Blood 91, 2240–2248 (1998).

40. Tsukuba, T. et al. Cathepsin E deficiency impairs autophagic proteolysis in macrophages. PLoS One 8, e82415 (2013).

41. Szulc-Dąbrowska, L., Bossowska-Nowicka, M., Struzik, J. & Toka, F.N. Cathepsins in Bacteria-Macrophage Interaction: Defenders or Victims of Circumstance? Front Cell Infect Microbiol 10, 601072 (2020).

42. Marion Chad, R., et al. Chitinase 3-Like 1 (Chil1) Regulates Survival and Macrophage-Mediated Interleukin-1β and Tumor Necrosis Factor Alpha during Pseudomonas aeruginosa Pneumonia. Infection and Immunity 84, 2094–2104 (2016).

43. Di Liberto, D., et al. Role of the chemokine decoy receptor D6 in balancing inflammation, immune activation, and antimicrobial resistance in Mycobacterium tuberculosis infection. Journal of Experimental Medicine 205, 2075–2084 (2008).

44. Castanheira, F.V.e.S., et al. The Atypical Chemokine Receptor ACKR2 is Protective Against Sepsis. Shock 49, 682–689 (2018).

45. Lima, I.L.d.A., et al. Contribution of atypical chemokine receptor 2/ackr2 in bone remodeling. Bone 101, 113–122 (2017).

46. Crosslin, D.R. et al. Genetic variation associated with circulating monocyte count in the eMERGE Network. Human Molecular Genetics 22, 2119–2127 (2013).

47. Sakaue, S. et al. A cross-population atlas of genetic associations for 220 human phenotypes. Nature Genetics 53, 1415–1424 (2021).

48. Kachuri, L. et al. Genetic determinants of blood-cell traits influence susceptibility to childhood acute lymphoblastic leukemia. The American Journal of Human Genetics 108, 1823–1835 (2021).

49. Vuckovic, D. et al. The Polygenic and Monogenic Basis of Blood Traits and Diseases. Cell 182, 1214–1231.e1211 (2020).

50. Astle, W.J. et al. The Allelic Landscape of Human Blood Cell Trait Variation and Links to Common Complex Disease. Cell 167, 1415–1429.e1419 (2016).

51. Liu, L. et al. Eosinophils attenuate arthritis by inducing M2 macrophage polarization via inhibiting the IκB/P38 MAPK signaling pathway. Biochemical and Biophysical Research Communications 508, 894–901 (2019).

52. Zhang, Y. et al. Eosinophils Reduce Chronic Inflammation in Adipose Tissue by Secreting Th2 Cytokines and Promoting M2 Macrophages Polarization. International Journal of Endocrinology 2015, 565760 (2015).

53. Wu, D. et al. Eosinophils sustain adipose alternatively activated macrophages associated with glucose homeostasis. Science 332, 243–247 (2011).

54. Shen, F.W. et al. Cloning of Ly-5 cDNA. Proceedings of the National Academy of Sciences 82, 7360–7363 (1985).

55. Jamieson, T. et al. The chemokine receptor D6 limits the inflammatory response in vivo. Nat Immunol 6, 403–411 (2005).

56. Cossarizza, A. et al. Guidelines for the use of flow cytometry and cell sorting in immunological studies (second edition). European Journal of Immunology 49, 1457–1973 (2019).

57. Wilson, G.J., Fukuoka, A., Vidler, F. & Graham, G.J. Diverse myeloid cells are recruited to the developing and inflamed mammary gland. Immunology 165, 206–218 (2022).

58. Kawamoto, T. & Kawamoto, K. Preparation of Thin Frozen Sections from Nonfixed and Undecalcified Hard Tissues Using Kawamoto’s Film Method (2020). Methods Mol Biol 2230, 259–281 (2021).

