## Supplementary Information for "Eosinophils promote monocyte to macrophage differentiation and anti-bacterial immunity"

### Supplementary Materials

### Methods

#### Cell isolation

Lavage of the pleural cavity space was carried out by injecting 2ml PBS containing 2mM EDTA.

#### Tissue Digestion

Mouse was shaved and an approx. 5 $cm^2$ piece of dorsal skin was excised and chopped finely, before digestion with HBSS containing collagenase D (1 mg/mL Roche), Dispase II (0.5 mg/mL, Roche) and DNase I (0.1 mg/mL, Roche). Cell suspensions were obtained from the spleen by filtering through a 70 µm EASYstrainer mesh (Greiner One). The omentum was removed and chopped finely before digestion with 2 mg/ml Collagenase type I (Roche) and 4% BSA (Invitrogen) in DMEM. Samples were shaken at 800 rpm, 37°C for 1 h. The omentum was then passed through a 40 µm EASYstrainer mesh (Greiner One). For each tissue, RBC were lysed in 1 mL RBC lysis buffer (Invitrogen) for 1 min, and washed with 10 ml PBS.

#### Macrophage depletion

150 µl Clophosome Liposomes containing either PBS or clodronate (Stratech) were injected intraperitoneally into mice on day 0. Mice were culled and peritoneal lavage was caried out after 1 and 4 weeks.

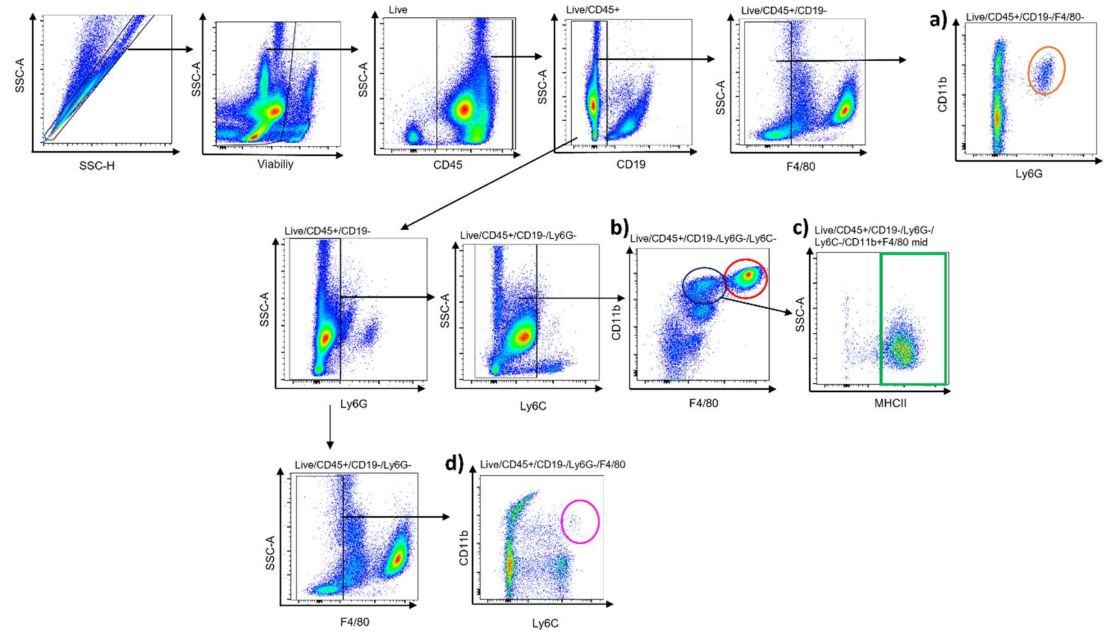


**Supplementary Figure 1: Gating strategy to define cell types in the peritoneal cavity.** Flow cytometry was carried out to identify cell types in the peritoneal cavity. Single cells were gated, dead cells were excluded, and CD45+ and CD19- cells were gated. a) Neutrophils were identified by further gating on F4/80- cells and defined as CD11b+, Ly6G+ (orange). **b)** Resident peritoneal macrophages, LPMs were defined as CD11b+, F4/80 high (red). **c)** Monocyte derived SPMs were further gated as F4/80 mid and MHCII positive (green). **d)** Monocytes were defined as Ly6G-, F4/80-, CD11b+, Ly6C+ (purple).

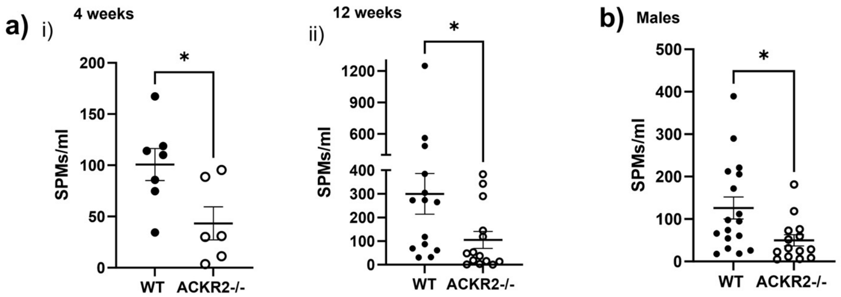


**Supplementary Figure 2: SPMs are reduced at different ages and in male mice.** **a)** Flow cytometry was carried out to determine the number of SPMs in WT (black circles) and ACKR2-/- (white circles) female **i)** 4 week old (WT, *n*=7, ACKR2-/-, *n*=8) and **ii)** 12 week old mice (*n*=14, per group). **b)** 8 week old males (WT, *n*=17, ACKR2-/-, *n*=14). Significantly different results are indicated (*, $p < 0.05$). Error bars represent SEM.

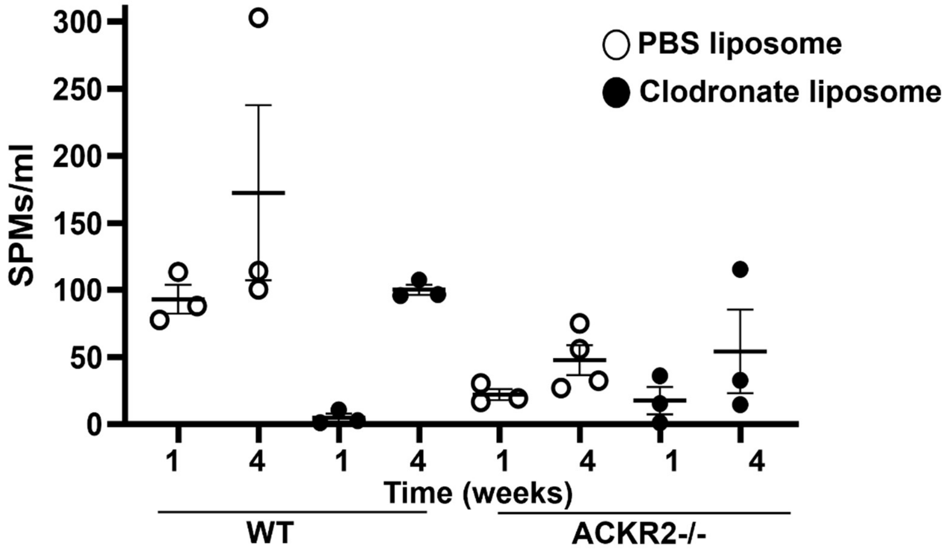


**Supplementary Figure 3: SPMs are replaced after clodronate liposome depletion. a)** Flow cytometry was carried out to determine the number of SPMs in the peritoneal cavity 1 and 4 weeks after i.p injection of 150 µl PBS (white circles) or clodronate (black circle) liposomes.

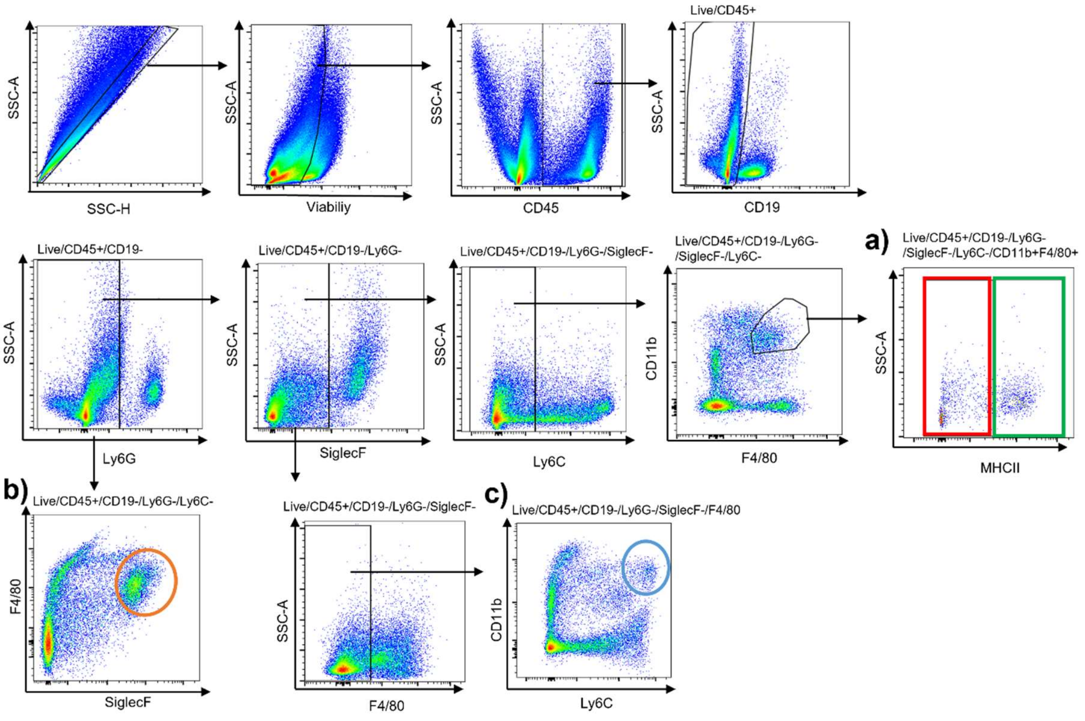


**Supplementary Figure 4: Gating strategy for cell types in the lung.** Flow cytometry was carried out to identify cell types in the lung. Single cells were gated, dead cells were excluded, and CD45+, CD19- cells and Ly6G- were gated. **a)** Interstitial macrophages were defined as SiglecF-, Ly6C-, CD11b+ and F4/80+, and further divided into MHCII negative (red) and positive (green) populations. **b)** Resident alveolar macrophages were defined as F4/80 low SiglecF+ (orange). **c)** Monocytes were defined as SiglecF-, F4/80-, CD11b+, Ly6C+ (blue).

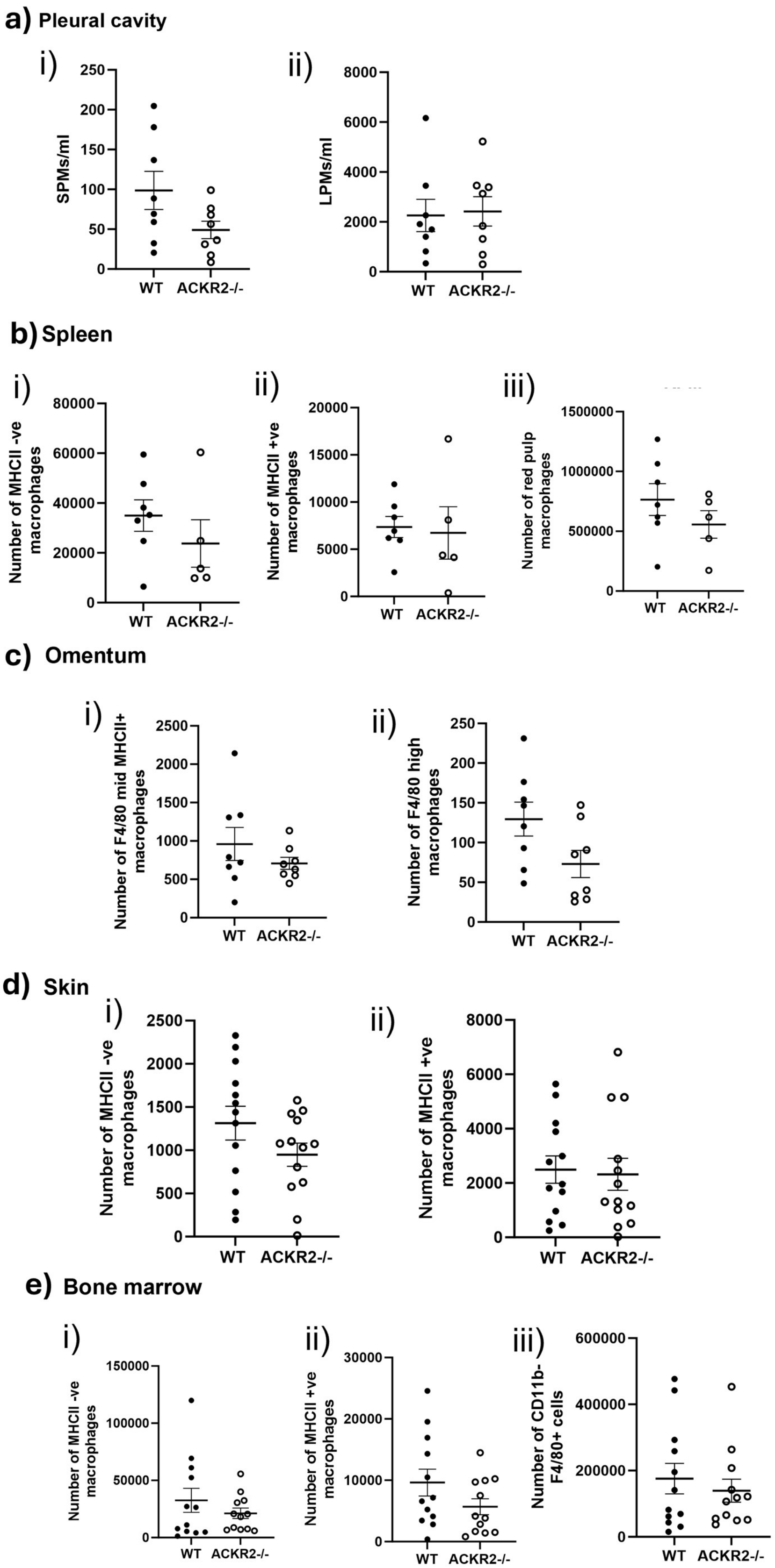
a) Pleural cavity
i)
SPMs/ml
250
200
150
100
50
0
WT
ACKR2-/-
ii)
LPMs/ml
8000
6000
4000
2000
0
WT
ACKR2-/-
b) Spleen
i)
Number of MHCII -ve macrophages
80000
60000
40000
20000
0
WT
ACKR2-/-
ii)
Number of MHCII +ve macrophages
20000
15000
10000
5000
0
WT
ACKR2-/-
iii)
Number of red pulp macrophages
1500000
1000000
500000
0
WT
ACKR2-/-
c) Omentum
i)
Number of F4/80 mid MHCII+ macrophages
2500
2000
1500
1000
500
0
WT
ACKR2-/-
ii)
Number of F4/80 high macrophages
250
200
150
100
50
0
WT
ACKR2-/-
d) Skin
i)
Number of MHCII -ve macrophages
2500
2000
1500
1000
500
0
WT
ACKR2-/-
ii)
Number of MHCII +ve macrophages
8000
6000
4000
2000
0
WT
ACKR2-/-
e) Bone marrow
i)
Number of MHCII -ve macrophages
150000
100000
50000
0
WT
ACKR2-/-
ii)
Number of MHCII +ve macrophages
30000
20000
10000
0
WT
ACKR2-/-
iii)
Number of CD11b- F4/80+ cells
600000
400000
200000
0
WT
ACKR2-/-

**Supplementary Figure 5: Macrophage numbers are not significantly altered in the absence of ACKR2 in other tissues within the body.** Flow cytometry was used to determine the number of macrophages within the single cell, live, CD45+, CD19-, Ly6G-, Ly6C- populations in various tissues. **a)** pleural cavity, **i)** SPMs, and **ii)** LPMs, WT, *n*=8 (black circles), ACKR2-/-, *n*=8 (white circles), were gated as in the peritoneal cavity (Supplementary Figure 1). **b)** Spleen macrophages were defined as CD11b+ F4/80+ and further divided into **i)** MHCII negative and **ii)** MHCII positive. **iii)** Resident red pulp macrophages were defined as CD11b-F4/80+, WT, *n*=7, ACKR2-/-, *n*=5. **c)** In the omentum macrophages were defined as **i)** CD11b+F4/80 mid MHCII positive, and **ii)** CD11b+F4/80 high, *n*=8, per group. **d)** Skin macrophages were defined as CD11b+ F4/80+ and further divided into **i)** MHCII negative and **ii)** MHCII positive, *n*=13 per group. **e)** Bone marrow macrophages were defined as CD11b+ F4/80+ **i)** MHCII negative and **ii)** MHCII positive, **iii)** CD11b-F4/80+, *n*=12, per group.

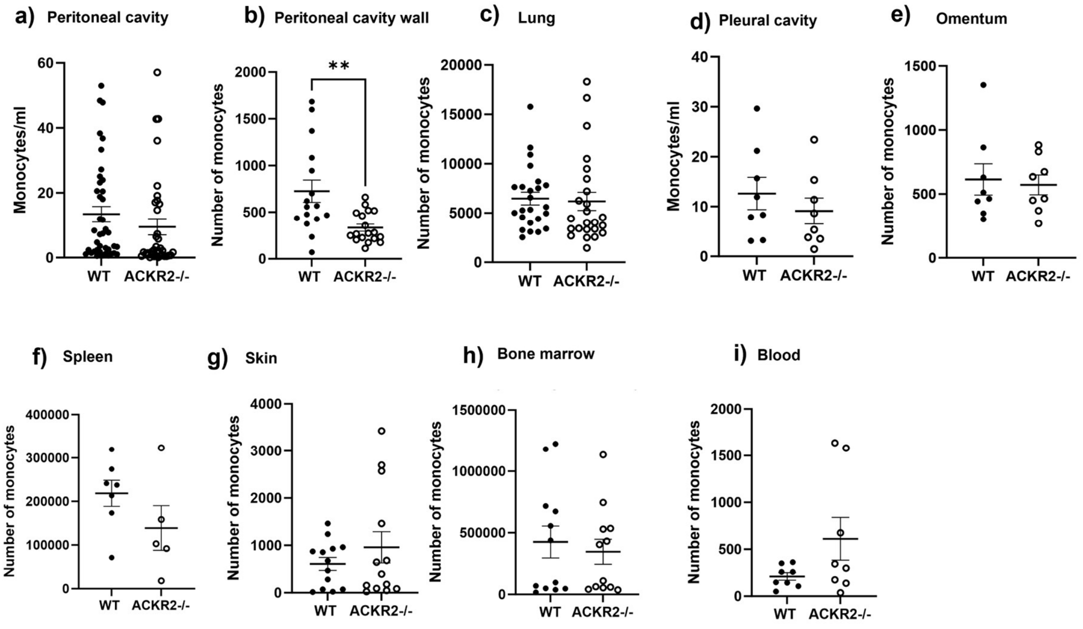


**Supplementary Figure 6: Monocyte numbers throughout the body.** Monocytes were defined by flow cytometry as single, live, CD45+, CD19-, Ly6G-, CD11b+ and Ly6C+ cells. Numbers were determined in WT (denoted by black circles) and ACKR2-/- (white circles) mice, in the **a)** peritoneal cavity (WT, *n*=40, ACKR2-/-, *n*=36), **b)** peritoneal cavity wall (WT, *n*=16, ACKR2-/-, *n*=18), **c)** lung (*n*=24, per group), **d)** pleural cavity, (*n*=8, per group) **e)** omentum (*n*=8, per group), **f)** spleen (WT, *n*=7, ACKR2-/-, *n*=5), **g)** skin (*n*=13, per group), **h)** bone marrow (*n*=12, per group), and **i)** blood (*n*=8, per group). Significantly different results are indicated (*, $p < 0.05$). Error bars represent SEM.

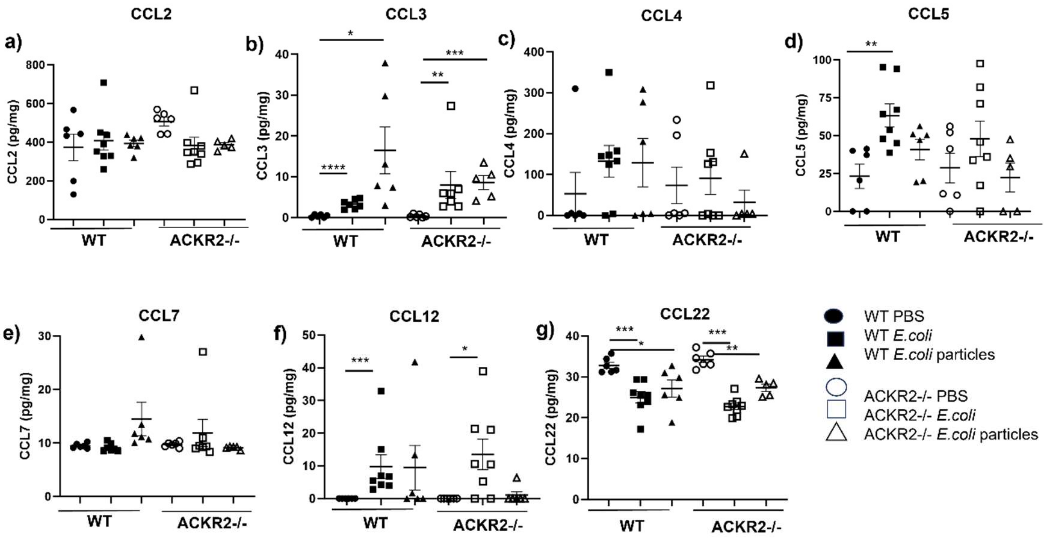


**Supplementary Figure 7: ACKR2 does not scavenge ligands in the peritoneal cavity during infection and inflammation.** Protein levels were determined by Luminex, after i.p challenge of WT (denoted by black shapes) and ACKR2-/- (white shapes) mice, with either PBS (circles), $5 \times 10^6$ CFU *E. coli* strain CFT073 (squares), or 200 µg *E.coli* particles (triangles) for 18 h (PBS, *n*=6 per group, WT *E.coli*, *n* = 8, WT *E.coli* particles, *n*= 6, ACKR2-/- *E.coli*, *n* = 8, ACKR2-/- *E.coli* particles, *n*= 5). Concentrations of the ACKR2 ligands, **a)** CCL2, **b)** CCL3, **c)** CCL4, **d)** CCL5, **e)** CCL7, **f)** CCL12, and **g)** CCL22, were determined. Significantly different results are indicated (*, $p < 0.05$). Error bars represent SEM.

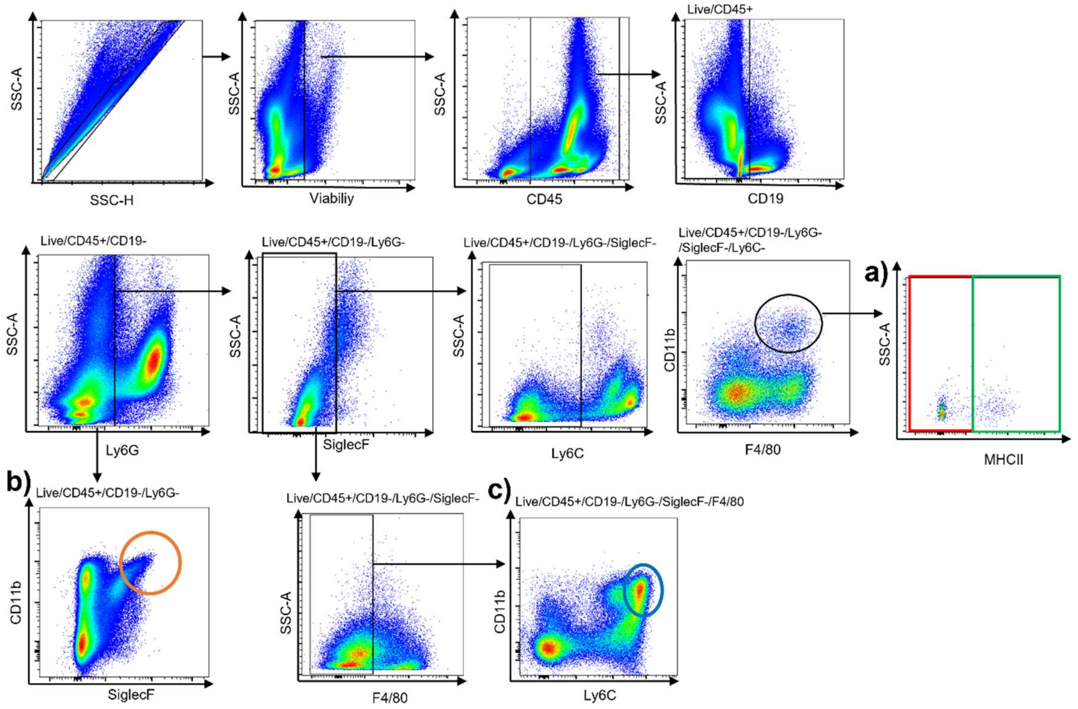


**Supplementary Figure 8: Gating strategy to define cell types in the bone marrow.** Flow cytometry was carried out to identify cell types in the bone marrow. Single cells were gated, dead cells were excluded, and CD45+, CD19-, and Ly6G- cells were gated. **a)** CD11b+ F4/80+ macrophages were further defined as MHCII negative (boxed in red) and MHCII positive (green). **b)** Eosinophils were gated as CD11b+ and SiglecF+ (orange). Monocytes were defined as Ly6G-, SiglecF-, F4/80-, CD11b+, Ly6C+ (blue).

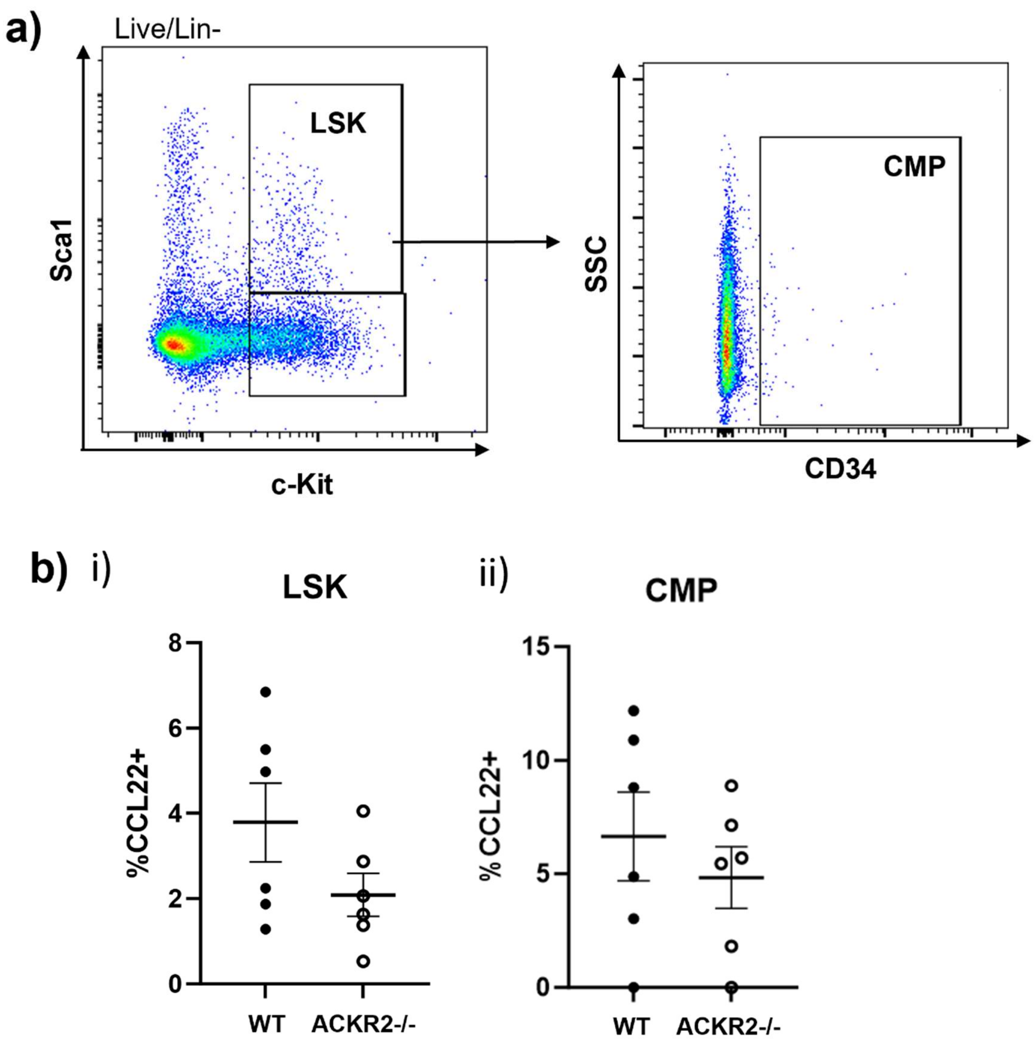


**Supplementary Figure 9: ACKR2 is not expressed by haematopoietic precursors in the bone marrow. a)** Flow cytometry gating to define LSK (Lin- Sca1+ c-kit+) cells, and common myeloid progenitors (CMP) within the bone marrow. **b)** ACKR2 expression was determined by comparing CCL22 uptake by **i)** LSK and **ii)** CMP from WT (black circles) and ACKR2-/- (white circles) mice.
